# Generally-healthy individuals with aberrant bowel movement frequencies show enrichment for microbially-derived blood metabolites associated with reduced kidney function

**DOI:** 10.1101/2023.03.04.531100

**Authors:** Johannes P. Johnson-Martínez, Christian Diener, Anne E. Levine, Tomasz Wilmanski, David L. Suskind, Alexandra Ralevski, Jennifer Hadlock, Andrew T. Magis, Leroy Hood, Noa Rappaport, Sean M. Gibbons

## Abstract

Bowel movement frequency (BMF) has been linked to changes in the composition of the human gut microbiome and to many chronic conditions, like metabolic disorders, neurodegenerative diseases, chronic kidney disease (CKD), and other intestinal pathologies like irritable bowel syndrome and inflammatory bowel disease. Lower BMF (constipation) can lead to compromised intestinal barrier integrity and a switch from saccharolytic to proteolytic fermentation within the microbiota, giving rise to microbially-derived toxins that may make their way into circulation and cause damage to organ systems. However, the connections between BMF, gut microbial metabolism, and the early-stage development and progression of chronic disease remain underexplored. Here, we examined the phenotypic impact of BMF variation in a cohort of generally-healthy, community dwelling adults with detailed clinical, lifestyle, and multi-omic data. We showed significant differences in microbially-derived blood plasma metabolites, gut bacterial genera, clinical chemistries, and lifestyle factors across BMF groups that have been linked to inflammation, cardiometabolic health, liver function, and CKD severity and progression. We found that the higher plasma levels of 3-indoxyl sulfate (3-IS), a microbially-derived metabolite associated with constipation, was in turn negatively associated with estimated glomerular filtration rate (eGFR), a measure of kidney function. Causal mediation analysis revealed that the effect of BMF on eGFR was significantly mediated by 3-IS. Finally, we identify self-reported diet, lifestyle, and psychological factors associated with BMF variation, which indicate several common-sense strategies for mitigating constipation and diarrhea. Overall, we suggest that aberrant BMF is an underappreciated risk factor in the development of chronic diseases, even in otherwise healthy populations.

## INTRODUCTION

The gut microbiome influences human health in a number of ways, from mediating early life immune system development ^1,2^, to determining personalized responses to nutritional interventions ^3,4^ and influencing the central nervous system ^5,6^. Stool transit time, defined as the rate at which stool moves through the gastrointestinal tract, is a major determinant of the composition of the human gut microbiota ^7^. Transit time is affected by diet, hydration, physical activity, host mucus production, microbe- and host-derived small molecules (e.g., short chain fatty acids, bile acids, or neurotransmitters), and peristaltic smooth muscle contractions in the gastrointestinal tract ^8,9^. Stool transit time can be partially estimated using the Bristol Stool Scale ^10^, edible dyes ^7^, indigestible food components (e.g., corn) ^11^, or self-reported bowel movement frequency (BMF) ^12,13^. Aberrant BMFs, in particular, have been implicated as risk factors in a number of chronic diseases ^14–16^.

Abnormally high BMF (e.g., diarrhea, defined as more than three watery stools per day), has been associated with lower gut microbiome alpha-diversity, inflammation, increased susceptibility to enteric pathogens, and poorer overall health ^12,17–19^. Abnormally low BMF (e.g. constipation, defined as fewer than three hard, dry stools per week), has been associated with higher gut microbiome alpha-diversity, reduced intestinal barrier integrity, enrichment in microbially-derived urinary metabolites known to be hepatotoxic or nephrotoxic, and with an increased risk for several chronic medical conditions, including neurodegenerative disorders and chronic kidney disease (CKD) ^14,20–22^. Indeed, constipation is a known risk factor for CKD severity and end-stage renal disease (ESRD) progression ^23,24^. In one study, up to 71% of dialysis patients suffered from constipation ^25^, while the prevalence of constipation in the general population was 14.5% in adults under 60 years old and 33.5% in those over 60 ^26^. A nationwide study of veterans found an incrementally higher risk for renal disease progression in those who reported increasingly severe constipation ^27^. However, while it is clear that morbidity and mortality risk worsen with constipation in those with active CKD, potential connections between BMF and the development and early-stage kidney disease are not yet established.

Both constipation and CKD associate with declines in gut microbiota-mediated short-chain fatty acid (SCFA) production and a rise in the production of amino acid putrefaction byproducts, including several toxic microbe-host co-metabolites, such as 3-indoxyl sulfate (3-IS), p-cresol sulfate (PCS) and phenylacetylglutamine (PAG), which all have been implicated in CKD progression ^28–30^. This is consistent with an established microbiota-wide transition from saccharolytic to proteolytic fermentation in constipated individuals due to the exhaustion of dietary fiber in stool ^14,31^. Thus, while the potential relationship between BMF and organ function in healthy populations is not fully understood, the gut metabolic phenotype associated with lower BMF suggests a connection.

In this study, we focus on categories of self-reported BMF in a large population of generally-healthy individuals with a wide range of molecular phenotypic data in order to quantify the phenotypic impact of BMF on blood plasma metabolites, blood proteins, clinical chemistries, and gut microbiome composition in a pre-disease context. By exploring the molecular phenotypic consequences of BMF variation in a generally-healthy cohort, along with BMF-associated demographic, dietary, lifestyle, and psychological factors, we aimed to identify early-stage biomarkers and potential therapeutic targets for the monitoring and prevention of certain chronic, non-communicable diseases, like CKD.

## RESULTS

### A cohort of generally-healthy individuals

3,955 Arivale Scientific Wellness program participants with BMF data were initially considered in this analysis. Arivale, Inc. (USA), was a consumer scientific wellness company that operated from 2015 until 2019. Briefly, participants consented to having their health, diet, and lifestyle surveyed through an extensive questionnaire, along with blood and stool sampling for multi-omic and blood plasma chemistries data generation (**Fig. 1**). Any respondents that indicated “true” or affirmatively to any of the following questionnaire features were excluded from the analysis (i.e., they were not considered “generally-healthy”): taking blood pressure, cholesterol, or laxative medication or having self or family history of bladder or kidney disease (i.e. kidney cancer, bladder infections, polycystic kidney disease or PKD, kidney stones, kidney failure or kidney disease), inflammatory bowel disease (IBD; both Crohn’s Disease and Ulcerative Colitis), irritable bowel syndrome (IBS), celiac disease, diverticulosis, gastroesophageal reflux disease (GERD), or peptic ulcers (i.e., these individuals were not considered ‘generally-healthy’–see Supplement, **Table S1**). There were 1,425 participants who met these exclusion criteria and had necessary covariate data. Across all Arivale participants that had available demographic and survey information, 82.8% of those individuals identified as “White” (N = 2,562), 8.5% identified as “Asian” (N = 262), 3.2% identified as “Black or African-American” (N = 98), 0.2% identified as “American Indian or Alaska Native” (N = 9), 0.65% identified as “Native Hawaiian or other Pacific Islander” (N = 20), and 4.7% identified as “Other” (N = 144). 93.6% of these individuals identified as “Non-Hispanic” (N = 2,897) and 6.4% identified as “Hispanic” (N = 198, 55.6% of which self-identify as “White”). Respondents were in the United States, predominantly from the Pacific West, and their ages ranged from 19 to 89 years old. 65.1% were female with a mean ± s.d. body mass index of 27.15 ± 5.89 (**Fig. S1**). 1,062 of these individuals had gut microbiome data, 486 had blood metabolomics data, 823 had proteomics data, 1,425 had clinical chemistries data, and 1,420 had survey data (derived from questionnaires). Self-reported BMF values (responses to typical number of bowel movements per week) were grouped into four categories (**Fig. 1**), which we labeled as: “constipation” (≤ 2 bowel movements per week), “low-normal” (3-6 bowel movements per week), “high-normal” (1-3 bowel movements per day), and “diarrhea” (4 or more bowel movements per day). We first looked at potential associations between BMF and relevant covariates: sex, age, BMI, estimated glomerular filtration rate (eGFR), low-density lipoprotein blood plasma levels (LDL), C-reactive protein blood plasma levels (CRP), hemoglobin A1c blood plasma levels (A1C), and the first three principal components of genetic ancestry (PC1, PC2, and PC3) (N = 1,425; **Fig. 2; Table S2**). When BMF was coded as an ordinal dependent variable and regressed using ordered proportional odds logistic regression (POLR), only BMI (POLR, FDR-corrected p = 1.82E-3), age (POLR, FDR-corrected, p = 2.07E-3), sex (POLR, FDR-corrected p = 3.68E-16), and the first three principal components of genetic ancestry (PC1, PC2, and PC3; POLR, FDR-corrected p < 0.0001) showed significant, independent associations with BMF (**Table S2**), with females, older individuals, and individuals with lower BMIs tending to report lower BMFs (**Fig. 2**). All covariates listed above were included in downstream regressions, regardless of whether or not they showed an independent association with BMF. The high-normal BMF group was chosen as the reference for all downstream regressions throughout the manuscript where BMF was encoded as a categorical variable. eGFR was also regressed against BMF and the other covariates to determine which were significant associated with eGFR, and the covariates with significant p-values included sex, age, BMI, LDL, A1C, PC1, PC2, and PC3 (GLM, p < 0.05).

**Figure 1.**
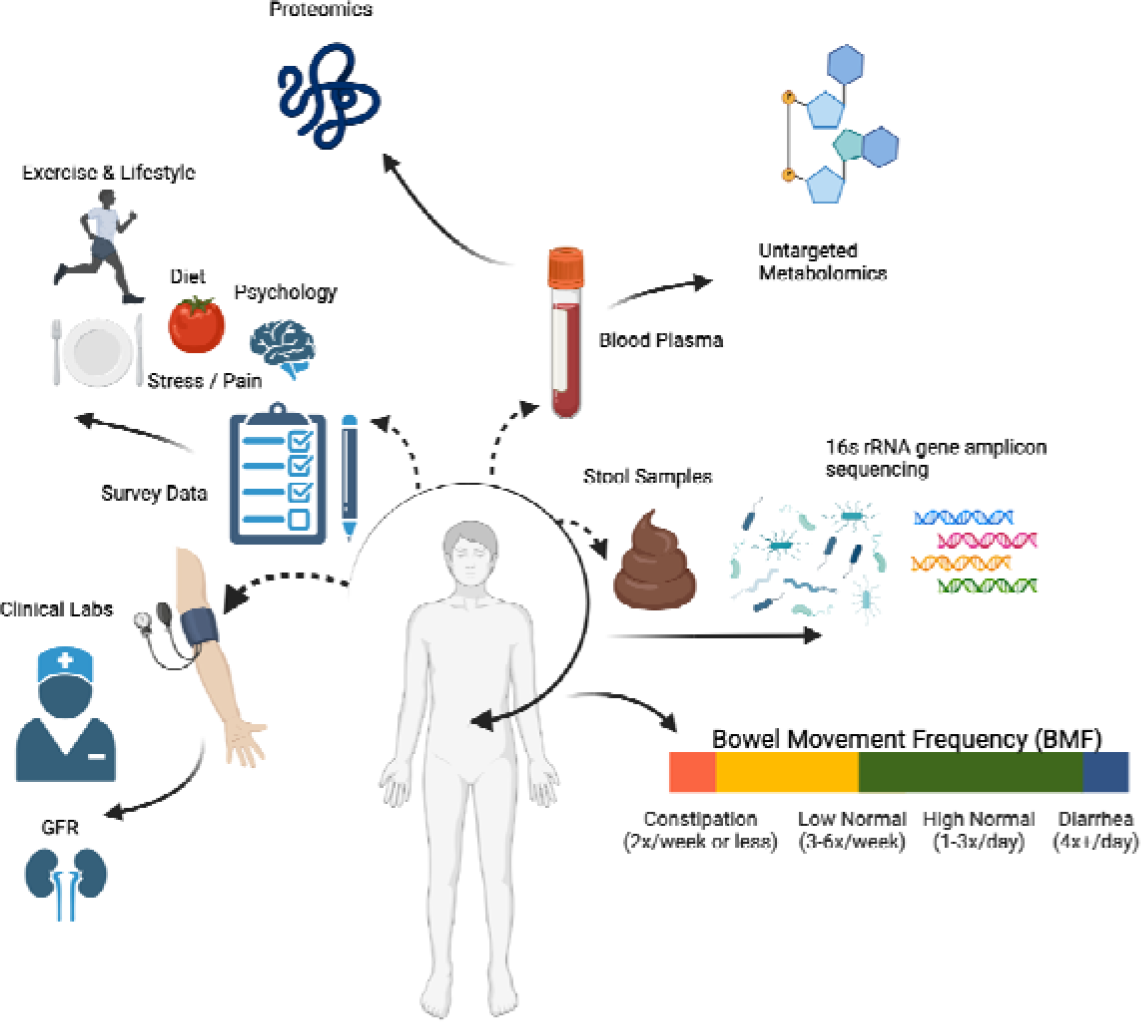
Data collection strategy. Arivale participants were sampled for blood plasma and stool, in addition to filling out extensive diet, health, and lifestyle questionnaires. Clinical chemistries, untargeted metabolomics, and proteomics data were generated from blood plasma samples. Gut microbiome 16S rRNA amplicon sequencing data were generated from stool samples collected using at-home kits. BMF data were extracted from the questionnaire data as self-reported frequencies per week or day.

**Figure 2.**
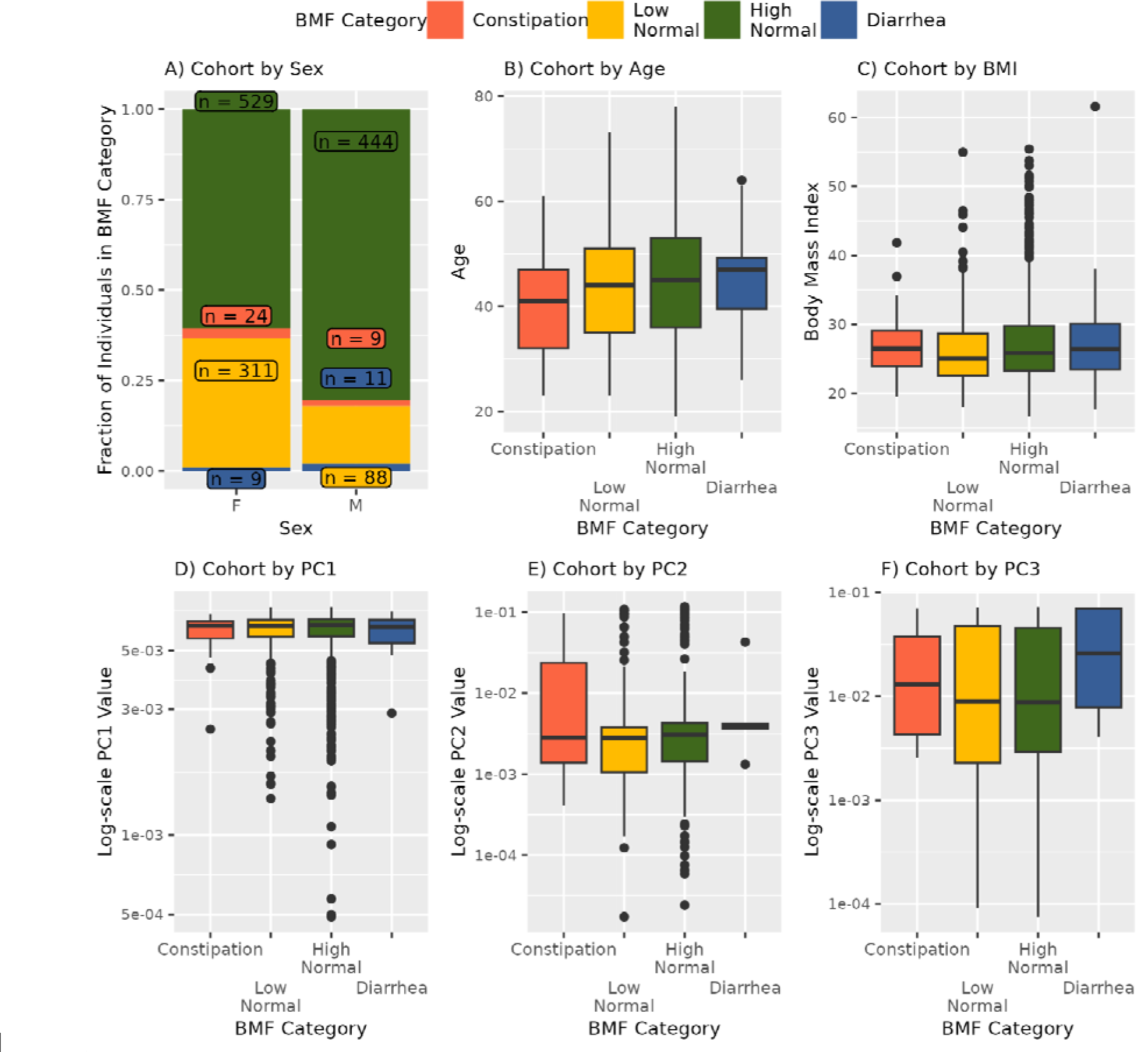
Plotting covariates that showed a significant association with BMF: sex, age, BMI, and the first three principal components of genetic ancestry (PC1-PC3) (A-F). POLR was used to regress BMF against the covariates (sex, age, BMI, eGFR, LDL, CRP, A1C, plus the first three principal components of genetic ancestry in the cohort, PC1, PC2, PC3). The result was that sex (p = 3.68E-16), BMI (p = 1.82E-3), age (p = 2.075E-3), and PCs1-3 (p < 0.00001, respectively) were significantly associated with BMF.

### Gut microbiome structure and composition across BMF categories

We looked at a subcohort of individuals that met our health exclusion criteria with 16S amplicon sequencing data from stool (N = 1,062). Amplicon sequence variant (ASV) richness (GLM, p = 2.85E-3, linear β_BMF_ = -65.9E-3) and Shannon diversity (GLM, p = 1.07E-3, linear β_BMF_= -3.25E-1) were negatively associated with BMF, independent of the covariates listed above, and with BMF encoded as an ordinal variable with a linear coefficient (**Fig. 3**). Pielou’s evenness, on the other hand, was positively associated with BMF (GLM, p = 8.5E-3, linear β_BMF_ = 2.6E-3), independent of covariates (**Fig. 3**).

**Figure 3.**
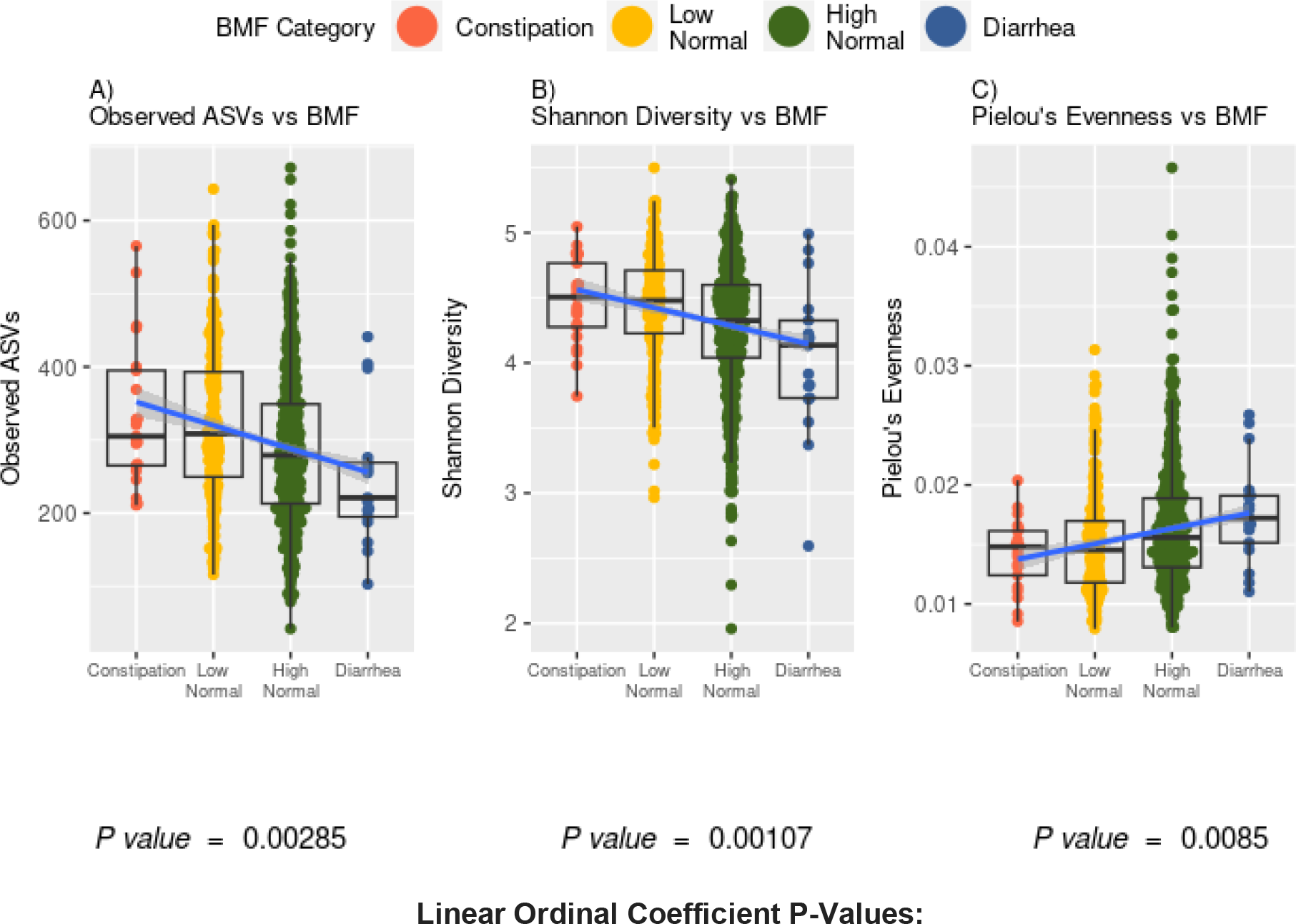
Associations between gut microbiome alpha-diversity and BMF. (A) Richness of amplicon sequence variants (ASVs) across BMF categories (ordinal BMF variable, Linear Regression, p = 2.85E-3). (B) Shannon Diversity across BMF categories (ordinal BMF variable, Linear Regression, p = 1.07E-3). (C) Pielou’s Evenness across BMF categories (ordinal BMF variable, Linear Regression, p = 8.5E-2).

Differential abundance analysis of commensal gut bacterial genera across BMF categories was conducted using beta-binomial regression (CORNCOB ^32^) with BMF encoded as a categorical variable with the high-normal group as the reference category. Of the 135 genera that passed our prevalence filter (i.e., detection across ≥ 30% of individuals), 59 were significantly associated with BMF (49 of which had genus-level taxonomic annotations; see **Table S1** for detailed list of β-coefficients and p-values), independent of covariates and following an FDR correction for multiple tests on the likelihood ratio test (LRT) p-values (FDR-corrected p < 0.05). We z-score normalized the centered log-ratio (CLR) abundances of the 49 annotated genera across all samples and then plotted the average z-score within each BMF bin for each taxon as a heatmap (**Fig. 4**). We also provide supplemental boxplots, showing CLR abundances across BMF categories, of the top 10 most abundant taxa and 10 taxa with the smallest p-values from the 49 mentioned above (**Fig. S2-S3**). In order of descending abundance, the following taxa were significantly enriched in constipated individuals, compared to the high-normal BMF category (Wald Test, FDR-corrected β_BMF_p < 0.05): *Ruminiclostridium_9, Ruminococcacaeae_UCG-005, Ruminococcaceae_NK41214_group, Family_XIII_AD3011_group, Romboutsia, Ruminocaccaeae_UCG-004, UBA1819, Negativibacillus, DTU089, GCA-900066225, Candidatus_Soleaferrea, Anaerotruncus, Defluviitaleaeceae_UCG-011, Eisenbergiella, Pygmalobacter, Peptococcus, Hydrogenoanaerobacterium, Anaerofustis*, and *DNF00809*. *Lachnospiraceae_ND3007_group* and *Lachnospiraceae_UCG-004* were significantly depleted in constipated individuals. Several more were associated with enrichment or depletion in the low-normal BMF category, compared to the reference category (**Fig. 4**; See Supplement). There was no significant difference between the high-normal and diarrhea categories for any of the genera, which could be due to low sample size in the diarrhea category (i.e., we were likely underpowered to detect those associations).

**Figure 4.**
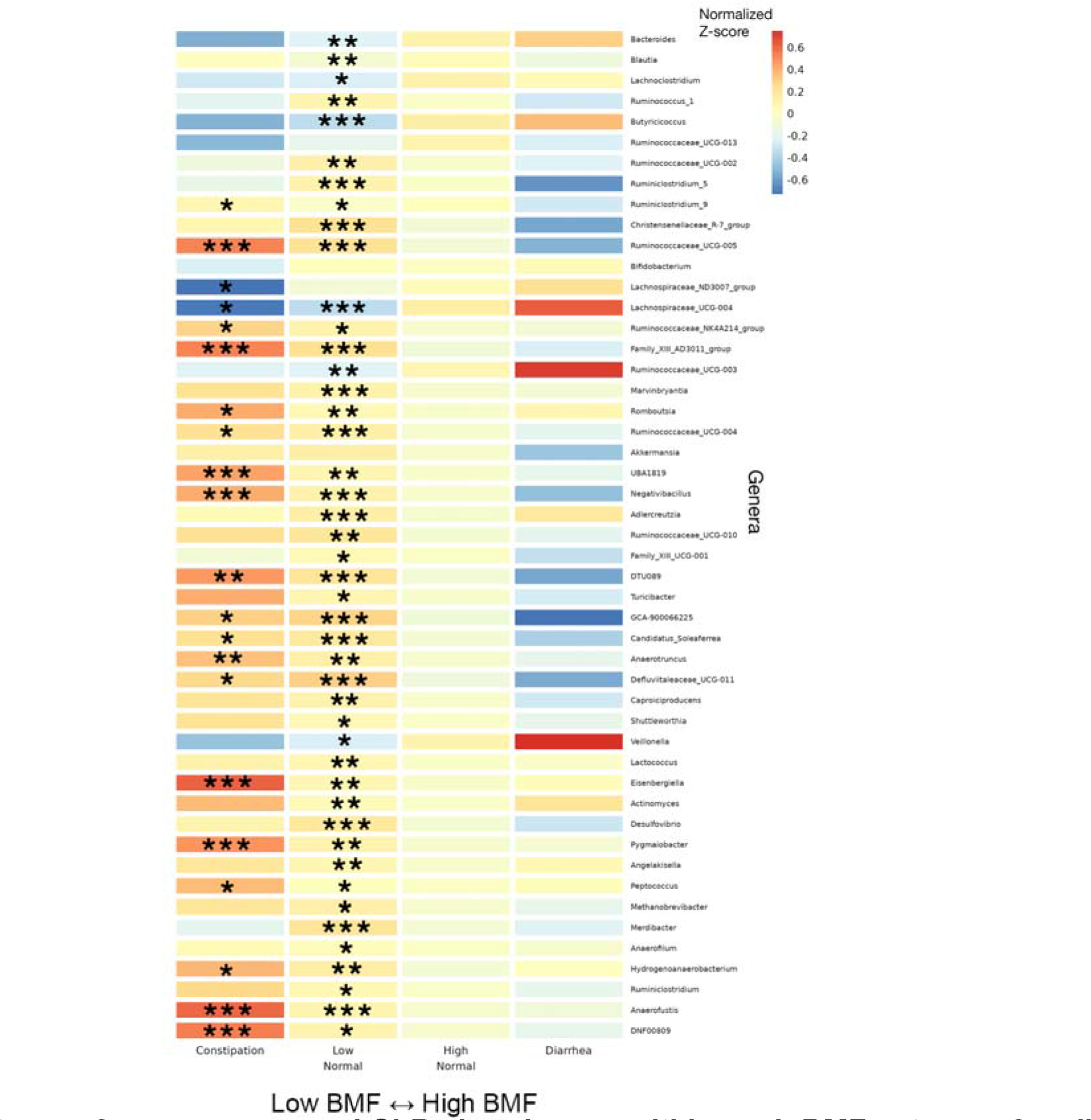
Heatmap of average z-scored CLR abundances within each BMF category for all annotated genera significantly associated with BMF. 46 significant taxa, in order of decreasing average relative abundance, with their z-scored, CLR-transformed abundances averaged within each BMF category plotted as a heatmap. Covariates included sex, age, BMI, eGFR, LDL, CRP, A1C, and PCs1-3. Asterisks denote the individual FDR-corrected significance threshold for the Wald Test p-value of the β_BMF_-coefficient for each BMF category, relative to the high-normal reference category. Rows without asterisks showed a significant overall model (FDR p-value <0.05), despite a lack of significance for the individual coefficients. (***): p < 0.0001, (**): 0.0001 < p < 0.01, (*): 0.01 < p < 0.05.

### Variation in blood metabolites across BMF categories

Blood metabolite vs. BMF regression analyses were run using a generalized linear modeling (GLM) framework in LIMMA, with BMF as a categorical independent variable, along with the same set of covariates mentioned above. Of the metabolites that passed our abundance and prevalence filters (N = 956, see **Method Details**), 9 unique metabolites were significantly associated with BMF (all 9 showed differential abundance between low-normal and high-normal categories, which is the comparison we were most powered for), independent of covariates and following an FDR correction for multiple tests (GLM, FDR-corrected p < 0.05, **Fig. 5**, **Table S2**). The annotated metabolites tended to show a decreasing trend with increasing BMF, while the unannotated metabolites and 3-IS showed more varied relationships (e.g. monotonic and non-monotonic) with BMF (**Fig. 5, S4**). PCS, PAG, PCG, and 3-IS were significantly enriched in the low-normal BMF category, compared to the reference category (**Fig. 5, S4**). 75 unique metabolites were significantly associated with eGFR, independent of covariates and following the same FDR correction for multiple tests (linear regression, FDR-corrected p < 0.05, **Fig. 5, S4; Table S4**). Only one of these eGFR-associated metabolites overlapped with any of the BMF-associated metabolites: 3-IS.

**Figure 5.**
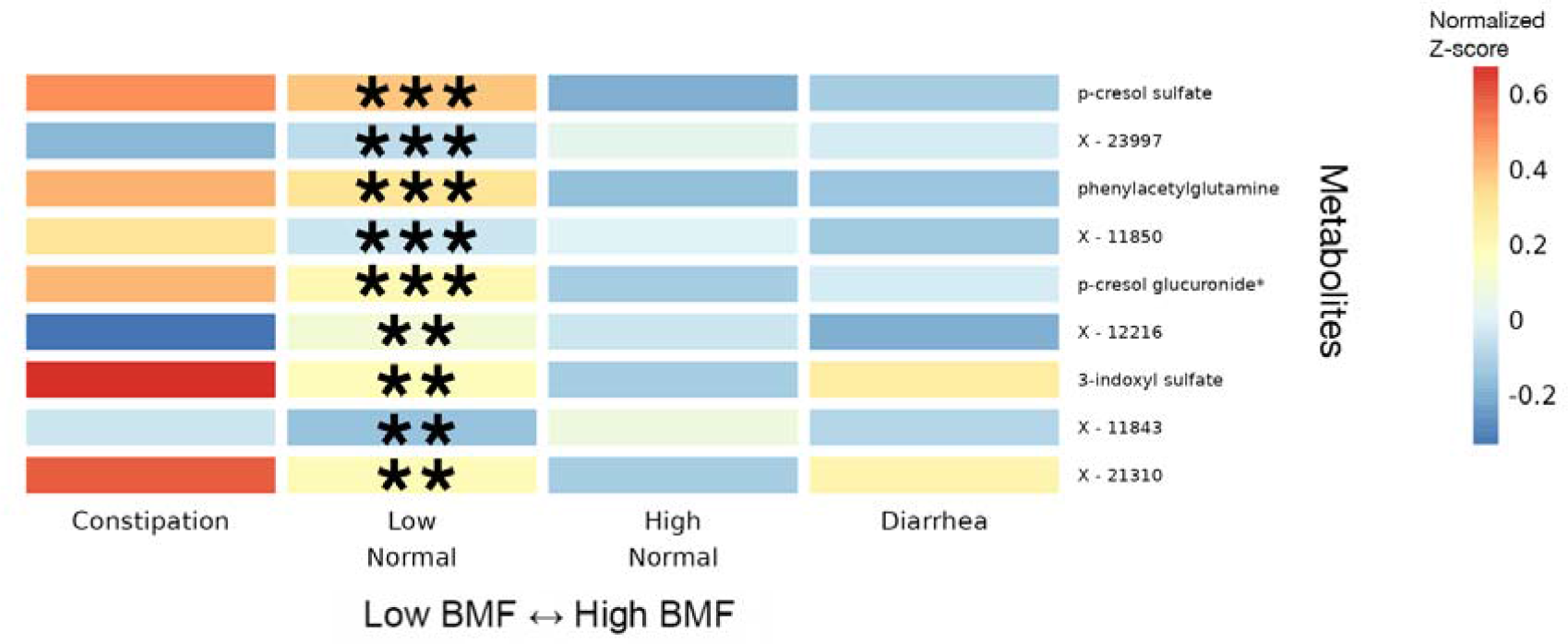
Heatmap of average z-scored blood plasma metabolites levels within each BMF category for all metabolites significantly associated with BMF. 11 significant blood plasma metabolites, with average z-scores within each BMF category plotted as a heatmap. Significant associations were identified using LIMMA, with FDR-corrected p-values of the ratio test between the main model and the null model. Here, the covariates included sex, age, BMI, eGFR, LDL, CRP, A1C, and PCs1-3. Asterisks denote metabolites with significant β_BMF_ coefficient(s) in the linear regression model after FDR correction. (***): p < 0.0001, (**): 0.0001 < p < 0.01, (*): 0.01 < p < 0.05.

### Blood plasma chemistries across BMF categories

Of the 55 blood plasma chemistries filtered for prevalence (see **Method Details**), 21 were significantly associated with diarrhea (e.g., omega-6 fatty acid, homocysteine, total protein, and bilirubin) and one (omega-6/omega-3 ratio in the blood) was associated with the low-normal BMF category, relative to the reference category, after adjusting for all covariates and for multiple testing (**Fig. 6**; N = 1,425, GLM, FDR-corrected p < 0.05).

**Figure 6.**
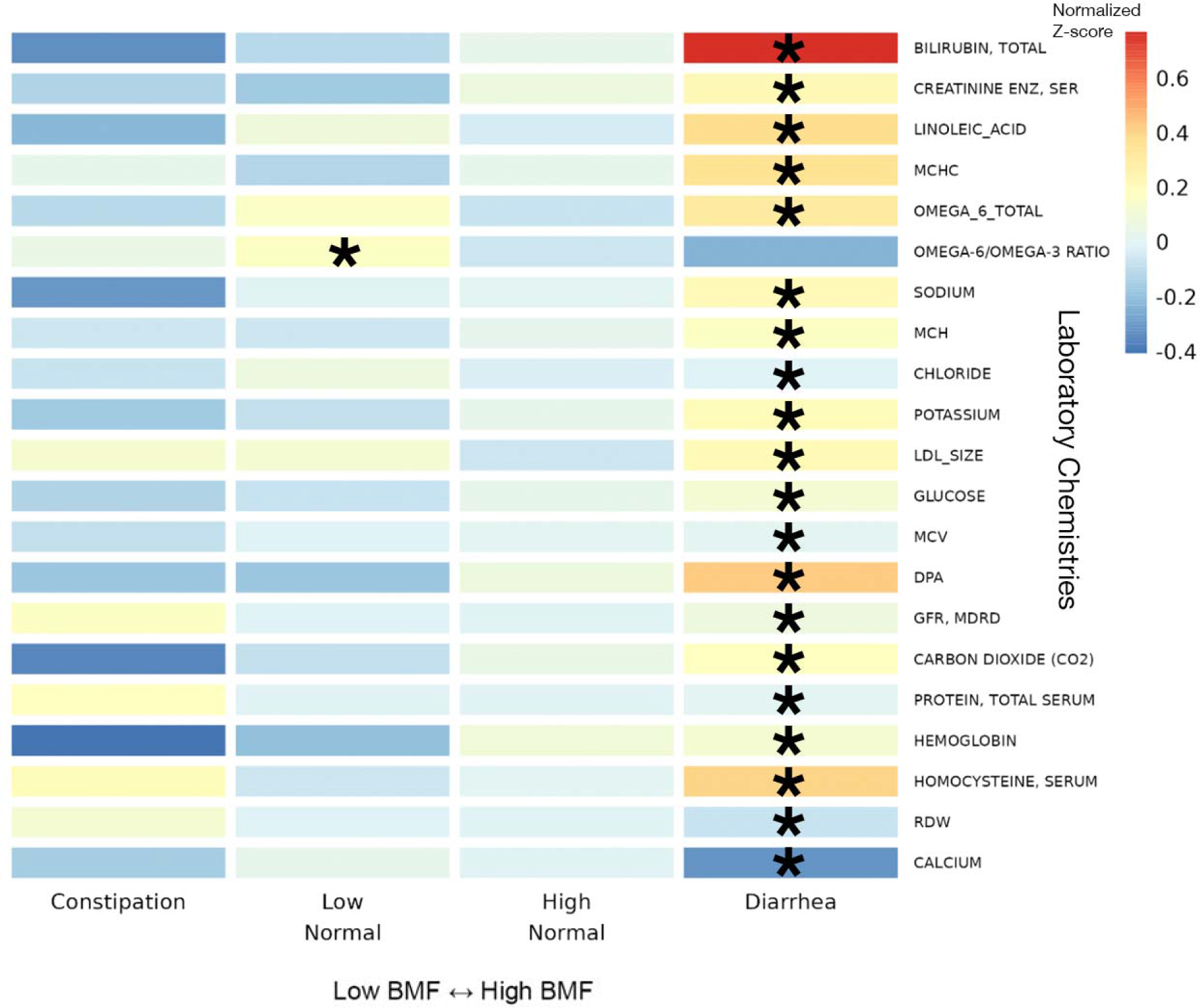
Heatmap of average z-scored clinical chemistries within each BMF category fo all chemistries significantly associated with BMF. 22 BMF-associated chemistries, identified using LIMMA models with FDR-corrected p-values of the ratio test between the main model and the null model, with average z-scores within each BMF category plotted as a heatmap. Here, the covariates included sex, age, BMI, eGFR, LDL, CRP, A1C, and PCs1-3. Asterisks denote FDR-corrected p-value thresholds for metabolites with significant β_BMF_ coefficient(s) in the linear regression model. (***): p < 0.0001, (**): 0.0001 < p < 0.01, (*): 0.01 < p < 0.05.

### Blood proteomics across BMF categories

None of the 274 blood proteins that passed our prevalence filter (see **Method Details**) showed significant associations with BMF after adjusting for all covariates and for multiple testing (N = 823, GLM, FDR-corrected p < 0.05).

### Self-reported diet, lifestyle, anxiety and depression histories associated with BMF categories and demographic covariates

99 survey questions (see **Supplement**; questions with sparse data were filtered out) on health, diet, and lifestyle were examined from 1,420 generally-healthy individuals from the Arivale cohort in order to identify covariate-independent associations with BMF. Tests were run using the “polr” package in R (ordinal regression) ^33^, including the same set of covariates from the prior analyses, and with BMF coded as a categorical variable with high-normal BMF as the reference group (**Fig. 7**). Response categories for each question ascended ordinally in value or intensity (i.e., low to high), so that a positive association represented an increase in that variable. Across the 99 questions, the top results with significant odds ratios related to BMF categories were displayed relative to high-normal BMF (**Fig. 7**), colored by the variable category (“Diet/Lifestyle” or “Health/Digestion”). BMI, age, sex, and other covariates were also associated with many of these questionnaire-derived features, independent of BMF (**Fig. 7**). In particular, females tended to eat more vegetables and fruit in a week and had a higher diarrhea frequency. Males, on the other hand, showed higher weekly snack intake and easier bowel movements (**Fig. 7**). Unsurprisingly, constipation (lowest BMF range) was negatively associated with reported ease of bowel movement and diarrhea was positively associated with self-reported diarrhea frequency (i.e., these were separate questions on the questionnaire) (**Fig. 7**). Those with higher weekly snack intake were more likely to be in the low-normal BMF category, and those with higher weekly vegetables intake, weekly fruit intake, greater ease of bowel movements, and those with higher self-reported diarrhea frequency were more likely to be in the high-normal BMF category (**Fig. 7**). Higher diarrhea frequency was significantly associated with having a higher BMI and with being younger relative to the rest of the cohort, while being older made one more likely to report having greater ease of bowel movement (**Fig. 7**). Finally, those with low LDL values (better cholesterol health) were more likely to report higher fruit intake and those with low CRP (low inflammation) values were more likely to report higher vegetables intake (**Fig. 7**). These findings showcase a variety of common-sense dietary and lifestyle factors that could be leveraged to manage BMF, cardiometabolic, and immune health.

**Figure 7.**
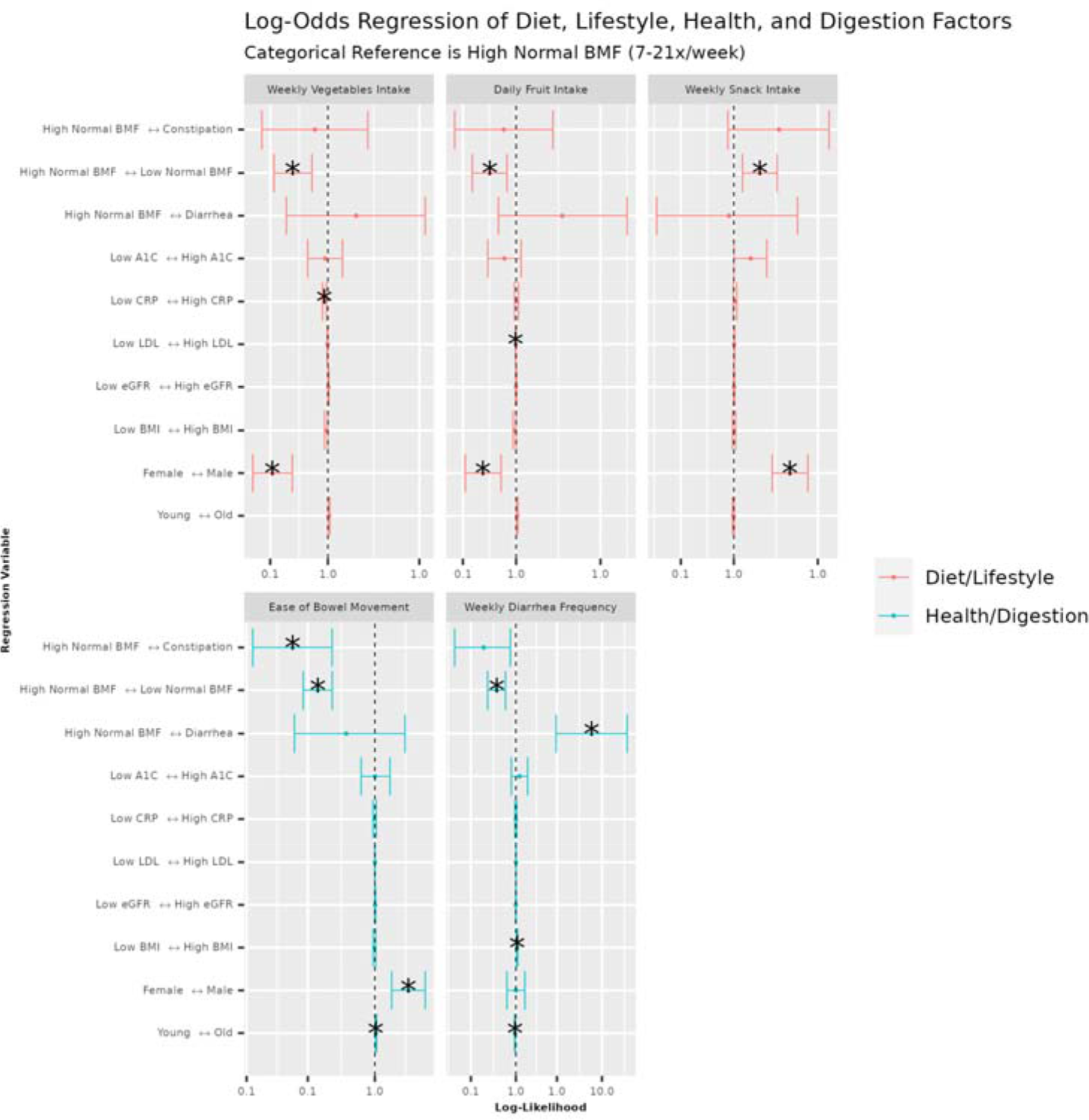
Ordinal regression odds ratio for health, diet, and lifestyle survey data vs BMF and covariates. Variables are colored by category: questions related to diet, exercise, and lifestyle (Diet/Lifestyle), and questions related to current digestive symptoms/function and health history (Health/Digestion). The BMF reference category was “high-normal” BMF (7-21 bowel movements per week). Each tick on the vertical axes represents a directional association in likelihood across the horizontal axis. The center line over the plots at x = 1.0 represents an equal likelihood of reporting an increase in number, intensity, frequency, or agreement (depending on the response variable) between the left side of the arrow on the vertical axis tick and the right side of the arrow on the vertical axis tick. A confidence interval that does not span the center line is significantly associated with the independent variable on the vertical axis tick. (*): FDR-corrected p-value < 0.05.

A subset of participants self-reported their history of depression and anxiety, including: “self-current”, “self-past”, and “family” history of depression and anxiety (N = 2,096, see Supplement; 11 questions related to anxiety and 23 related to depression). After logistic regression, 3 “true or false”-response questions related to history of depression in self and family history appeared marginally significant (logistic regression, FDR-corrected p < 0.1), with a self-reported “true” response to a “family history of depression” showing a marginal association with constipation (logistic regression, FDR-corrected < 0.1), a self-reported “true” response to a “sibling history of depression” showing a significant association with diarrhea (logistic regression, FDR-corrected < 0.05), and a self-reported “true” response to “recent ailments; self-history of depression” showing a marginal association with low-normal BMF (logistic regression, FDR-corrected < 0.1). Similarly, the same approach yielded a single marginal association between a “true” response to “self past history of anxiety disorder” and low-normal BMF (logistic regression, FDR-corrected < 0.1). Each of these associations were relative to the high-normal BMF reference category.

*BMF-associated blood metabolites associated with kidney function in a generally-healthy cohort* Using the nine BMF-associated metabolites (ordered in ascending p-value: PCS, X - 23997, PAG, X - 11850, PCG, X - 12216, 3-IS, X - 11843, and X - 21310), an analysis was performed on all of the generally-healthy Arivale participants with paired BMF, eGFR, and blood metabolomic data (N = 572). Using OLS, eGFR was regressed against BMF (encoded as a numerical variable between 1, 2, 3, or 4, with 1 being constipation, 2 being low-normal, 3 being high-normal, and 4 being diarrhea) and the nine BMF-related metabolites, which yielded a significant overall model (**Fig. S8**; OLS, R^2^ = 0.082, p = 2.42E-7). Two of the BMF-associated metabolites showed significant beta-coefficients in the model: X - 12216 and 3-IS (**Fig. S8**; OLS, β_X-12216_ = -1.98, p = 1.20E-2 and β_3-IS_ = -9.69, p = 1.96E-5, respectively). These negative coefficients indicated that higher baseline levels of these blood metabolites were associated with lower kidney function.

Finally, given that microbially-derived 3-IS was independently associated with both eGFR and BMF, we hypothesized that 3-IS may be mediating, in part, the impact of BMF on eGFR. To test this hypothesis, we ran a causal mediation analysis (using the mediation library in R ^34^; see **Methods**) on the generally-healthy Arivale individuals with BMF, eGFR, and the blood metabolomics data (N = 572; **Fig. 8; S7**). BMF categories were merged into a “Low” (low-normal BMF and constipation) and a “High” categories (high-normal BMF and diarrhea participants) in order to consolidate the BMF categories with very small Ns (i.e., constipation and diarrhea). The total effect of the overall model did not quite pass our significance threshold of alpha < 0.05 (total effect, p = 0.064), but we saw a significant average direct effect of BMF on eGFR (ADE = -4.458, p = 0.012) and a highly significant average causal mediation effect of BMF via 3-IS on eGFR (ACME = 1.343, p < 2E-16; **Fig. 8**). A similar analysis was performed on those respondents that had vegetables intake data, and a marginally significant average direct effect (ADE, p = 0.058) and total effect (p = 0.062) were observed for an outcome model of eGFR ∼ 3-IS + vegetables intake (merged into a “Low” and “High” category, with “High” being the control value) + BMF (merged into a “Low” and “High” category) and a mediation model of 3- IS ∼ vegetables intake (merged) and BMF (merged).

**Figure 8.**
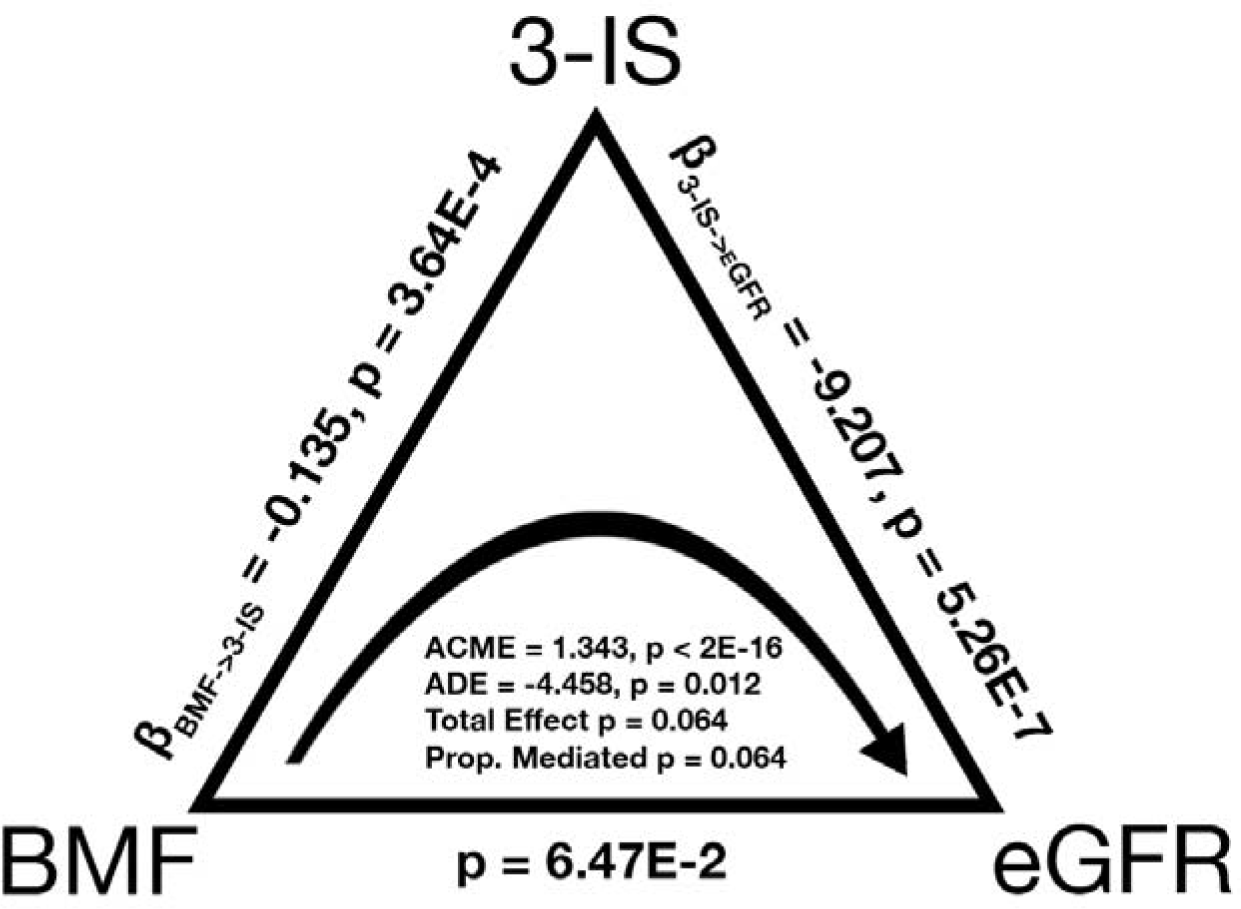
Causal mediation analysis, with BMF as the treatment variable, 3-IS as the mediator variable, and eGFR as the response variable. **The average direct effect (ADE) of** BMF on eGFR and the average causal mediated effect (ACME) of BMF on eGFR via 3-IS were found to be significant (N = 572; ADE -4.458, p = 0.012; ACME 1.343 p < 2E-16). The total effect and the proportion mediated terms did not pass our significance threshold of alpha=0.05.

## DISCUSSION

In this study, we delve into the multi-omic fingerprint of cross-sectional BMF variation in a large, generally-healthy population (**Fig. 1**). We find that aberrant BMFs were associated with variation in the ecological composition of the gut microbiota, plasma metabolite levels, clinical chemistries, diet, lifestyle, and psychological factors (**Figs. 4-7**). Overall, we observe an enrichment of microbially-derived uremic toxins in blood resulting from protein fermentation in the guts of individuals with lower BMFs. These toxins have been implicated in disease progression and mortality in CKD ^24,35^ and many of the same metabolites have been associated with other chronic diseases, like neurodegeneration ^36,37^.

Of the core set of covariates used in our regression analyses, only age, sex, BMI, and genetic ancestry PCs 1-3 were independently associated with BMF, with females, individuals with lower BMIs, and younger individuals showing lower average BMFs (**Fig. 2**). Consistent with these results, women are known to be at higher risk of constipation and kidney dysfunction ^38,39^. In a prior study, individuals with lower BMIs were shown to produce less motilin (i.e., a hormone involved in gut motility) and were more likely to suffer from constipation ^40^. Lower BMFs have also been linked to inflammation, oxidative stress, and cardiovascular disease risk ^41,42^. The associations between BMF and the first three principal components of genetic ancestry indicate a relationship between host genetics and BMF variation, which is further supported by a prior GWAS study ^43^.

Independent of these covariates, several gut bacterial genera enriched in individuals with lower BMFs (CORNCOB, p < 0.001), such as *Christensenellaceae_R-7_group*, *Anaerotruncus*, *Blautia*, *Family_XIII_AD3011_group* (Anaerovoracaceae family), and *Methanobrevibacter*, were previously found to be enriched in Parkinson’s disease (PD) patients who often suffer from chronic constipation ^44^*. Desulfovibrio*, which has been shown to be enriched in several disease states ^45^, was elevated at lower BMF (**Fig. 4**). Another set of genera were depleted in lower BMF categories, such as *Bacteroides*, *Lachnoclostridium*, *Lachnospiraceae_ND3007_group*, *Lachnospiraceae_UCG-004*, and *Veillonella*, which are all important contributors to SCFA production ^46–49^. This reduction in SCFA producers is consistent with the switch away from saccharolytic fermentation towards proteolytic fermentation in the case of constipation ^14^. Reduced SCFA production is known to weaken smooth muscle contractions that drive peristalsis ^50–52^, acting as a positive feedback on constipation.

Furthermore, constipation can induce mechanical damage to the gut epithelium ^53–55^, which may in turn contribute to higher systemic inflammation and disruptions to epithelial integrity ^35,56,57^. This epithelial damage, combined with chronic inflammation, may allow for excess luminal metabolites to leak into the bloodstream, including toxic protein fermentation byproducts, which could cause tissue damage throughout the body and exacerbate conditions like CKD ^35,58–60^.

Consistent with our microbiome results, we found gut microbiome-derived protein fermentation byproducts, like PCS, PAG, and 3-IS, were enriched in the blood of individuals with lower BMFs (**Fig. 5**) ^61–63^ . PCS has been associated with deteriorating kidney function and with damage to nephrons as well as cognitive decline and neuroinflammation ^64,65^. 3-IS has been associated with vascular disease and mortality in CKD patients ^66^. PAG has been associated with CKD progression and mortality ^29,30,61,62^ . Ultimately, we see an enrichment in microbially-derived uremic toxins in the blood of generally-healthy individuals with lower BMFs.

Most of the clinical chemistry-BMF associations showed relative enrichment in the higher-BMF category, and these features tended to reflect hepatic and nephrotic function. For example, high bilirubin can indicate liver disease from the overactive breakdown of red blood cells, but interestingly, higher bilirubin levels in serum coincide with a lower risk for CKD development and progression, which coincides with our observation that the lowest BMF categories had higher levels of uremic toxins but lower bilirubin levels ^67^. Other metrics, like creatinine levels and linoleic acid levels, correlate positively with BMF and negatively with kidney function ^68–70^. In fact, most of the laboratory values, such as the mean corpuscular hemoglobin concentration (MCHC), which measures the concentration of blood cells, can indicate kidney or liver disease ^71^. It is interesting to note that biomarkers indicating kidney disease risk and progression were enriched at lower BMFs and biomarkers indicating liver disease risk and progression were enriched at higher BMFs in a generally-healthy population, showing how aberrant BMF in either direction may increase chronic disease risk.

In addition to demographic factors associated with BMF, the questionnaire results indicate dietary and lifestyle factors that are known to influence BMF, like fruit and vegetable intake (i.e., sources of dietary fiber and polyphenols) ^39,41^. We observed a lower fruit and vegetable intake and an increased likelihood of snacking in the low-normal BMF category compared to the high-normal BMF category ^26,39^. We also found that constipation and diarrhea were marginally (and in one case, significantly) associated with self-reported measures of depression and anxiety, which aligns with prior work showing higher prevalence of anxiety and depression (between 22-33%) on the Hospital Anxiety and Depression Scale (HADS) and the Mini International Neuropsychiatric Interview (MINI) in patients with chronic constipation ^72^.

Blood levels of 3-IS were independently associated with both BMF and eGFR, which led us to the hypothesis that 3-IS may mediate the potential influence of BMF on eGFR. Indeed, we observed a significant average direct effect of BMF on eGFR (ADE, p = 0.012) and a highly significant average causal mediation effect for 3-IS (ACME, p < 2E-16; **Fig. 8**). Together, these results indicate that aberrant BMF-associated increases in 3-IS are associated with declining kidney function in a generally-healthy cohort, which is consistent with similar associations that have been observed between 3-IS and poorer outcomes in CKD patients ^66^.

Bowel movement abnormalities, such as constipation or diarrhea, have been linked to diseases ranging from enteric infections ^19^ to many chronic diseases like CKD, IBD, and neurodegenerative conditions like Alzheimer’s and PD ^36,73,74^. Indeed, even in the context of our generally-healthy cohort, we see the build up of microbially-derived uremic toxins in the blood of individuals with lower BMFs. Perhaps most concerning was our observation that aberrant BMF- associated microbial metabolite 3-IS was also associated with lower eGFR values. In conclusion, we suggest that chronic constipation or diarrhea may be underappreciated drivers of organ damage and chronic disease, even in healthy populations. Our results underscore common-sense dietary and lifestyle changes, like increasing intake of fruits and vegetables, which may help to normalize BMF and perhaps reduce BMF-associated chronic disease risk.

## Study Limitations

There are some important limitations to consider when interpreting the results of this study. The generally-healthy cohort studied here was overwhelmingly “White”, predominantly female, and from the West Coast of the U.S.A., which limits the generalizability of our results. In addition, the diet, lifestyle, and mood data were self-reported and subject to biases and errors, and are not indicative of clinical diagnoses, although BMF was binned into four coarse-grained categories in an attempt to mitigate self-reporting bias. In fact, BMF is not quite synonymous with transit time through the gut, which can be measured through means like the “blue dye method” for transit time ^7^ , although BMF still appears to be a useful measure of self-reported bowel habit differences in this study when binned in such coarse-grained categories. We had limited representation in the constipation and diarrhea categories, which reflects the “generally-healthy” nature of this cohort, but this also limited our statistical power for detecting associations in these groups. The dietary variables that were associated with better BMF outcomes (i.e., increased dietary fiber intake, in the form of fruits and vegetables) are not devoid of clinical risk and may not be appropriate for everyone. For example, high-fiber diets can sometimes lead to bloating and inflammation in IBD patients ^75^. Additionally, CKD patients are often coached to limit their intake of fiber-rich foods because they can contain high levels of potassium and phosphorus ^76^. However, low-fiber diets may act as a positive feedback on constipation and inflammation. This highlights the importance of intervening at the prodromal stage, before disease manifests, when a healthy, plant-based diet is well-tolerated. While we find some evidence for microbially- derived, BMF-associated uremic toxins in blood influencing kidney function in a generally- healthy cohort, more work is needed to establish a link between longer-term BMF management and chronic disease risk. In addition, for the mediation analysis, we did not see a strong intervention effect or total model effect, despite seeing a highly significant mediation effect. This kind of result is expected when the treatment effect and the mediation effect are similar in magnitude, when there are opposing effect directions between treatments and mediators, or when there are other more complicated effects (e.g., non-linear associations) ^77^. Ultimately, future intervention trials should be done to assess the potential for managing BMF throughout the lifespan as a strategy to reduce chronic disease risk.

## ACKNOWLEDGMENTS

We thank Amy Willis for helpful advice on ordinal regression and for members of the Gibbons, Hood-Price, and Hadlock labs for helpful discussions on this work. This research was funded by Washington Research Foundation Distinguished Investigator Award and by startup funds from the Institute for Systems Biology (to S.M.G.). Research reported in this publication was supported by the National Institute of Diabetes and Digestive and Kidney Diseases of the National Institutes of Health (NIH) under award no. R01DK133468 (to S.M.G.) and by a Global Grants for Gut Health Award from Nature Portfolio and Yakult (to S.M.G.). The content is solely the responsibility of the authors and does not necessarily represent the official views of the NIH. The funders had no role in designing, carrying out or interpreting the work presented in the manuscript.

## AUTHORS CONTRIBUTIONS

J.P.J. and S.M.G. conceived of the study. J.P.J. conducted the analyses, wrote the code, and wrote the first draft of the manuscript. S.M.G. provided supervision. C.D., T.W., A.E.L., and A.R. contributed code and expert input on the analyses. A.E.L., T.W., D.L.S., A.R., J.H., A.T.M., L.H., and N.R. contributed to interpretation of the results and to editing the final manuscript.

## DECLARATION OF INTERESTS

L.H. is a former shareholder of Arivale. A.T.M. was a former employee of Arivale. Arivale is no longer a commercially operating company as of April 2019. The remaining authors report no competing interests.

## STAR METHODS

### Resource Availability

#### Lead Contact

Additional requests and information regarding resources, experimental materials, reagents, and assay vendors should be directed to and will be fulfilled by the lead contact, Sean Gibbons.

#### Materials Availability

This study did not generate new unique reagents.

#### Data and Code Availability

● Code used to analyze 16S rRNA gene amplicon sequencing data can be found at https://github.com/gibbons-lab/mbtools. Code used to run the statistical analyses described in this paper is available at https://github.com/jajohnso29/Generally-Healthy-Cohort-BMF.
● Qualified researchers can access the full Arivale deidentified dataset, including all raw data, supporting the findings in this study for research purposes through signing a Data Use Agreement (DUA). Inquiries to access the data can be made at and will be responded to within 7 business days.

### EXPERIMENTAL MODEL AND STUDY PARTICIPANT DETAILS

#### Institutional review board approval for the study

The procedures for this study were reviewed and approved by the Western Institutional Review Board, under the institutional review board study number 20170658 for the Institute for Systems Biology and 1178906 for Arivale, Inc.

#### Generally-healthy cohort

All study participants were subscribers in the Arivale Scientific Wellness program (2015-2019) and provided informed consent for the use of their anonymized, de-identified data for research purposes. Participants were community-dwelling, residents of Washington State and California (which are slightly leaner and healthier than other parts of the USA), over the age of 19, non-pregnant, but were not screened for the presence or absence of any particular disease. Participants provided detailed questionnaire data that included self-reported information about medical conditions and medications, along with blood and stool samples that were used to generate blood plasma metabolomics, proteomics, chemistries, and gut microbiome data (**Fig 1** and **Table S1**).

Only baseline time point samples were used for each participant for the baseline ‘omics analyses. A 30% prevalence filter was implemented across the gut microbiome, blood plasma metabolomics, proteomics, chemistries, and ordinal questionnaire data analyses. This meant that each final feature in the data could contain no more than 70% missing data from the final cohort of samples in order to be retained for downstream analysis. For microbiome analyses, a filtered subcohort of 1,062 individuals with ASV-level taxa counts, BMF, sex, age, eGFR, BMI, LDL, CRP, A1C, and genetic ancestry data were selected. This filtering resulted in a total of 135 genera. For the metabolomics analysis, a cohort of 486 participants with BMF, sex, age, eGFR, BMI, CRP, LDL, A1C, PC1, PC2, and PC3, and blood metabolomics data were selected. 956 metabolites were retained for downstream analyses. 274 blood proteins that met the prevalence (≥ 50%) filter in the cohort of 823 individuals were retained for downstream analyses. A ≥ 30% prevalence filter was applied to yield 1,425 samples with blood plasma clinical laboratory chemistries data, resulting in 55 features retained for downstream analyses. Similarly, for ordinal regression of the questionnaire data (e.g. diet, lifestyle, and stress/pain/health factors,) using the respective R package, polr ^33^, we collated all the responses and filtered out questions that contained more than 10% “NAs” (≥ 90% prevalence; and for binary variables in downstream depression/anxiety analyses: ≥ 10% affirmative or “True” responses). We also excluded binary response variables for the general survey questionnaire analysis (separate from the anxiety/depression analysis, which leveraged binary response features), which are incompatible with ordinal regression, resulting in 138 variables across 1,420 participants, in addition to having paired data on age, sex, eGFR, BMI, BMF, CRP, LDL, A1C, PC1, PC2, and PC3. The final features considered needed to retain at least 2 non-missing factors (or categories) and contain at least 10 responses per category, which resulted in 99 features. BMF data was captured from responses to a survey question on how many bowel movements an individual has per week, on average. The available responses to this question were: (1) Twice per week or less; (2) 3-6 times per week; (3) 1-3 times daily; or (4) 4 or more times daily. While the normal range of BMF encompasses both the second and third responses to this question (i.e., between three times a week and three times a day) ^78^, we chose to define 1-3 times per day (high-normal) as the reference group for the purposes of regression.

Finally, we imposed disease-related exclusion criteria, in order to generate a “generally-healthy” sub-cohort. These include any participants who reported affirmative or “true” to a history of taking cholesterol, laxative, or blood pressure medication, as well as those who reported a self- or family- history presence of the following diseases: bladder or kidney disease, inflammatory bowel disease (IBD), celiac disease, diverticulosis, gastroesophageal reflux disease (GERD), irritable bowel syndrome (IBS), or peptic ulcers (See **Fig. S1** in Supplement). 988 (25%) out of the initial 3,955 Arivale individuals with BMF data were excluded by these filters.

### METHOD DETAILS

#### Gut Microbiome Data

Fecal samples from Arivale participants were collected (described in Diener et al ^12^ and detailed here) from proprietary at-home kits developed by two microbiome vendors (DNA Genotek and Second Genome). Using the KingFisher Flex instrument, the MoBio PowerMag Soil DNA isolation kit (QIAGEN) enabled the isolation of stool DNA from 250 ml of homogenized human feces, after performing an additional glass bead-beating step. Qubit measurement and spectrophotometry were also performed using an A260/A280 absorbance ratio.

16S amplicon sequencing was run on a MiSeq (Illumina, USA) with either paired-end 300-bp protocol (DNA Genotek) or paired-end 250-bp protocol (SecondGenome). The FASTQ files were provided by the Illumina Basespace platform after the phiX reads were removed with basecalling. Length cutoffs of 250-bp for the forward reads and 230-bp for the reverse reads were employed. Any reads with more than 2 expected errors or ambiguous base calls under the Illumina error model were eliminated. Over 97% of the reads passed these filters, resulting in approximately 200,000 reads per sample.

Final truncated and filtered reads were then used to infer amplicon sequence variants (ASVs) with DADA2 ^79^. Each sequencing run separately resulted in its own error profiles. The final ASVs and counts were then joined, with chimeras removed using DADA2’s “consensus” strategy. After this step, ∼16% of reads were removed. Taxonomic assignment of ASVs was then achieved using the naive Bayes classifier in DADA2 with the SILVA database (version 128)^80^.

Nearly 90% of the ASVs were classified down to the genus level, which was the taxonomic level chosen for this analysis. 3,694 samples across 609 taxa were available from these methods, which were then filtered down to 135 taxa after using a 30% prevalence filter (no more than 70% of data was permitted to be missing per filtered taxa). Samples were rarefied to an even depth of 13,703 reads prior to calculating alpha-diversity metrics (using the “rarefy_even_depth( )” function in the phyloseq R package ^81^; rng seed = 111). ASV richness (Observed ASVs), Shannon Diversity, and Pielou’s evenness were calculated. Merging with covariate data resulted in 1,062 samples with 135 taxa for downstream analyses.

#### Olink Proteomics

Blood plasma proteomic data were generated by Olink Biosciences using the ProSeek Cardiovascular II, Cardiovascular III, and Inflammation arrays. The proteins were filtered down to 274 proteins and 823 samples, retaining proteins with ≥ 50% prevalence across samples and samples with the full set of covariate data. Post-filtering, NAN values were assumed to be below detection and imputed to be the median across samples for that particular protein. The values used for the proteomics analysis were from protein readings previously batch-corrected and normalized based on the overlapping reference samples within the batch plates (i.e., a set of Arivale plasma samples that are run with each batch). The corrected values were also scale- shifted to the reference sample and the original delivered data (using the seventh run as a baseline). Olink’s Proximity Extension Assay (PEA), a 2-antibody-barcode technology, is used to tag protein biomarkers with a proximity probe (which binds specifically to the target protein biomarker) and an extension probe (which carries a unique DNA barcode sequence) as described by Illumina in conjunction with Olink ^82–84^. Once both probes bind to each other due to a protein-protein interaction or by proximity, they trigger the activation of the extension probe, beginning the hybridization of the probe with a detection bead’s complementary DNA sequence. Each bead contains an individual identifier, which allows target proteins to be decoded according to a barcode. These methods are also described further in Zubair et al ^85^.

#### Metabolon Metabolomics

Metabolon obtained metabolomics data on the previously mentioned plasma samples using preparation, quality control, and collection methods described in previous studies ^86^. During sample processing, the plasma samples were thawed and proteins were removed using methanol extraction. Samples were then divided into 5 fractions including a backup fraction. Organic solvents were removed using TurboVap and measurements were then performed using high-performance liquid chromatography (HPLC) and high-resolution mass-spectrometry (MS). Four separate measurements were performed using different fractions combinations: positive-ion and negative-ion modes optimized for both hydrophobic and hydrophilic compounds. Batch correction was performed using quality control samples (i.e., a set of Arivale plasma samples that were run with each batch) and abundance data were normalized to these quality control samples. Metabolites were annotated according to 3 standards: Tier 1, matching to an internal standard; Tier 2, matching to a published MS spectrum; or Tier 3, matching to a known chemical formula. Unknown metabolites were unannotated and labeled with an “X - “ label followed by a unique identifier ^87^. 956 total metabolites showed at least 70% prevalence across 486 samples. In this analysis, missing values were imputed to be the median of the non-missing samples for each metabolite, and final downstream metabolites were log-transformed and merged with the full set of covariates.

For the multi-linear regression and causal mediation analysis, those with paired eGFR, BMF-associated metabolomics results, and BMF were filtered using the “generally-healthy” exclusionary criteria and the previously mentioned prevalence filtering for metabolomics. The remaining individuals (**Fig. 8,S7**; N = 572) were processed in a multi-linear regression (OLS) with eGFR ∼ BMF (encoded as a value between 1 and 4 with 4 being diarrhea or the highest BMF) + the obtained metabolomics values for the 9 BMF-associated metabolites (**Fig. S7-S8).** The other multi-omics covariates (sex, age, BMI, CRP, LDL, A1C, and PC1-PC3) were not considered for the subsequent mediation analysis (**Fig. 8**; N = 562), which was performed using a mediation model with the mediate( ) function from the mediation package in R ^88^. Using this modeling function, the outcome model was specified as eGFR ∼ 3-IS + BMF (where BMF was encoded as a binary categorical variable, with “Low” including those with low-normal BMF and constipation, and “High” containing those with high-normal BMF and diarrhea. “Low” was the control value for BMF and “High” was the treatment value) and the mediation model was assumed to be 3-IS ∼ BMF. ACME and ADE values were obtained from the model and reported using the diagram in **Fig. 8**. A GLM was also performed between eGFR ∼ BMF, 3-IS ∼ BMF, and eGFR ∼ 3-IS to obtain the β-coefficients and p-values for the relationships between the mediated variables (**Fig. 8**). Ultimately, we also performed a similar mediation analysis as before, but with the outcome model including eGFR ∼ 3-IS + BMF + vegetables intake and a mediation model containing 3-IS regressed against BMF + vegetables intake. This modeling strategy was applied to those with questionnaire survey data (N = 571) on vegetable eating habits (respondents claiming to eat 1 or less vegetables per day were in the “Low” treatment group, while those eating more vegetables than that daily were in the “High” control group) for the participants that self-responded to the inquiry of daily vegetable eating habits, implying a relationship between dieting factors and BMF on eGFR values through the proxy of 3-IS.

#### Blood Plasma Chemistries

LabCorp and Quest phlebotomists collected blood from Arivale participants using methods described previously by Wilmanski et al and others ^12^. Individuals were asked to abstain from alcohol, vigorous exercise, monosodium glutamate and aspartame at least 24 hours prior to drawing of the blood, as well as fasting at least 12 hours beforehand. Blood samples were collected for clinical chemistries, metabolomics and proteomics at the same time, and within 21 days of stool sampling. BMI was calculated from weight and height using the following formula 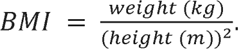 4,881 samples and 68 laboratory values were filtered down using the same prevalence filtering as the metabolomics data. 1,425 samples and 55 chemistries were retained. The final 55 features were log-transformed, with missing samples imputed to be the median value of the non-missing samples. These features were merged with the full set of covariates. eGFR was calculated based on the CKD Epidemiology Collaboration (CKD-EPI) creatinine Equation, as recommended by the current guidelines of the National Kidney Foundation ^89^: eGFR = 142 ✕ min(Scr/κ, 1)^α^ ✕ max(Scr/κ, 1)^-1.200^ ✕ 0.9938^Age^ ✕ 1.012 [if female], where Scr = standardized serum creatinine in mg/dL, κ = 0.7 (female) or 0.9 (male), and α = -0.241 (female) or -0.302 (male).

#### Questionnaire Data

3,482 self-reported questionnaire features were retrieved across 5,764 Arivale participants. After health and prevalence filtration, 138 downstream features remained, which were subsequently filtered down again to 99 final features by removing factored features with fewer than 10 responses per level and keeping features with at least 2 non-missing levels to the factor. Category responses were organized and numbered to be ordinally ascending in magnitude or intensity, with relatively even-spaced differences in magnitude between categories wherever possible (i.e. for a factored feature with levels from 1,…,n, the level labeled “1” represented responses such as “Strongly Disagree”, “Never”, “None”, or the lowest frequency/intensity, and the level labeled “n” represented responses such as “Strongly Agree”, “Always”, or the greatest frequency or intensity). These features were merged with the full set of covariate data.

#### Depression and Anxiety Health History Data

We used logistic regression to scrutinize associations between 23 (anxiety) and 35 (depression) independent binary (“true” or “false”) self-reported questions based on “self-current”, “self-past”, and “family” histories of depression or anxiety, with depression or anxiety encoded as a binary dependent variable, and BMF encoded as a categorical independent variable, and with the standard set of covariates.

### QUANTIFICATION AND STATISTICAL ANALYSIS

#### Statistical Analyses

The response variables were either: centered log ratio-transformed bacterial genus data, log-transformed plasma metabolomics data, batch-corrected plasma proteomics data, log-transformed plasma chemistries data, or ordinal response variables from questionnaire data, depending on the analysis. For the blood proteomics, plasma chemistries, and metabolite associations, generalized linear regression models were run using the LIMMA package in R ^90^. BMF was encoded as a categorical variable (or in the case of analyzing alpha-diversity, it was also computed as an ordinal variable with a linear model coefficient) with categories: 1 = constipation (1-2 bowel movements per week), 2 = low-normal (3-6 bowel movements per week), 3 = high-normal (1-3 bowel movements per day), and 4 = diarrhea (4 or more bowel movements per day). To begin characterizing the main variables in the cohorts: BMF and eGFR, a POLR regression (N = 1,425) was performed on BMF (encoded as an ordinal variable with categories “Constipation”, “Low Normal”, “High Normal”, and “Diarrhea” BMF in ascending order of magnitude) ∼ eGFR + other covariates (sex, age, BMI, CRP, LDL, A1C, PC1, PC2, and PC3). Similarly, a GLM (N = 1,425) was computed for eGFR ∼ BMF (also encoded ordinally) + other covariates (sex, age, BMI, CRP, LDL, A1C, PC1, PC2, and PC3). These were used to determine the significant covariates affecting each subsequent analysis (**Fig. 2**). Next, in each baseline regression, the following covariates were all included: sex, age, BMI, eGFR, CRP, LDL, A1C, PC1, PC2, and PC3. Gut bacterial genus-level counts were modeled with a beta-binomial distribution using the CORNCOB package in R ^32^. For the questionnaire data (ordinal response categories across diet, exercise, stress, pain, and other lifestyle factors), polr in R was used for the ordinal regression analysis (POLR). For the anxiety and depression data, which were binary in response (“True” or “False”; Non-responders to each feature were not considered and features were filtered to have at least 5 non-missing responses for each binary outcome), logistic regression was performed using the “glm(family = “binomial”)” function in R. All questionnaire and anxiety/depression response modeling results were FDR-corrected for significance. Finally, for the Arivale cohort, the initial time point or baseline value for eGFR was obtained alongside the initial or earliest time point sample for the BMF-related metabolites. eGFR was regressed against the BMF-associated metabolites in an OLS-based linear regression to determine visible effects of these metabolites on our available samples. Finally, a mediation analysis was run using the mediate( ) function in the mediation library available for R ^34^ on the individuals who met the “generally-healthy” exclusion criteria with paired eGFR, BMF, and 3-IS data. BMF was the treatment variable, 3-IS was the mediator, and eGFR was the response variable. ACME, ADE, total effect and proportion mediated were determined with nonparametric bootstrap confidence intervals.

## SUPPLEMENTAL FIGURES

**Figure S1.**
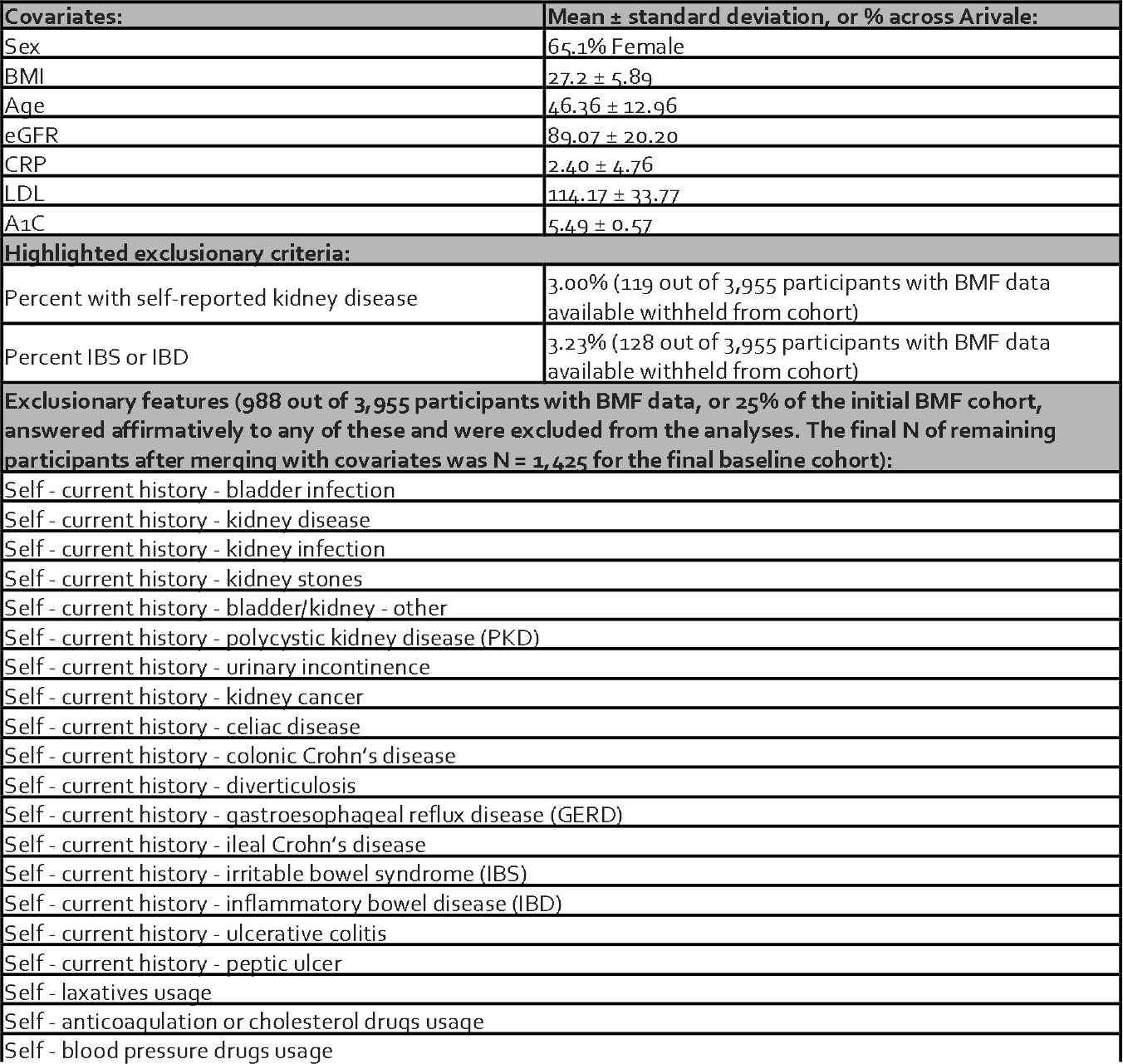
The modeling covariates and exclusionary criteria. Out of the 3,955 total Arivale participants that had BMF data, 3.00% self-reported kidney disease (the kidney-related questions in the exclusionary features) and 3.23% self-reported IBS or IBD. An initial baseline cohort of 3,132 participants that had health history survey questionnaire data was available. The participants that answered affirmatively to the exclusionary features were removed from the analysis, resulting in 25% of the initial cohort with BMF data being filtered down to N = 1,561, and subsequently, a final baseline cohort of 1,425 individuals after merging for covariates.

**Figure S2.**
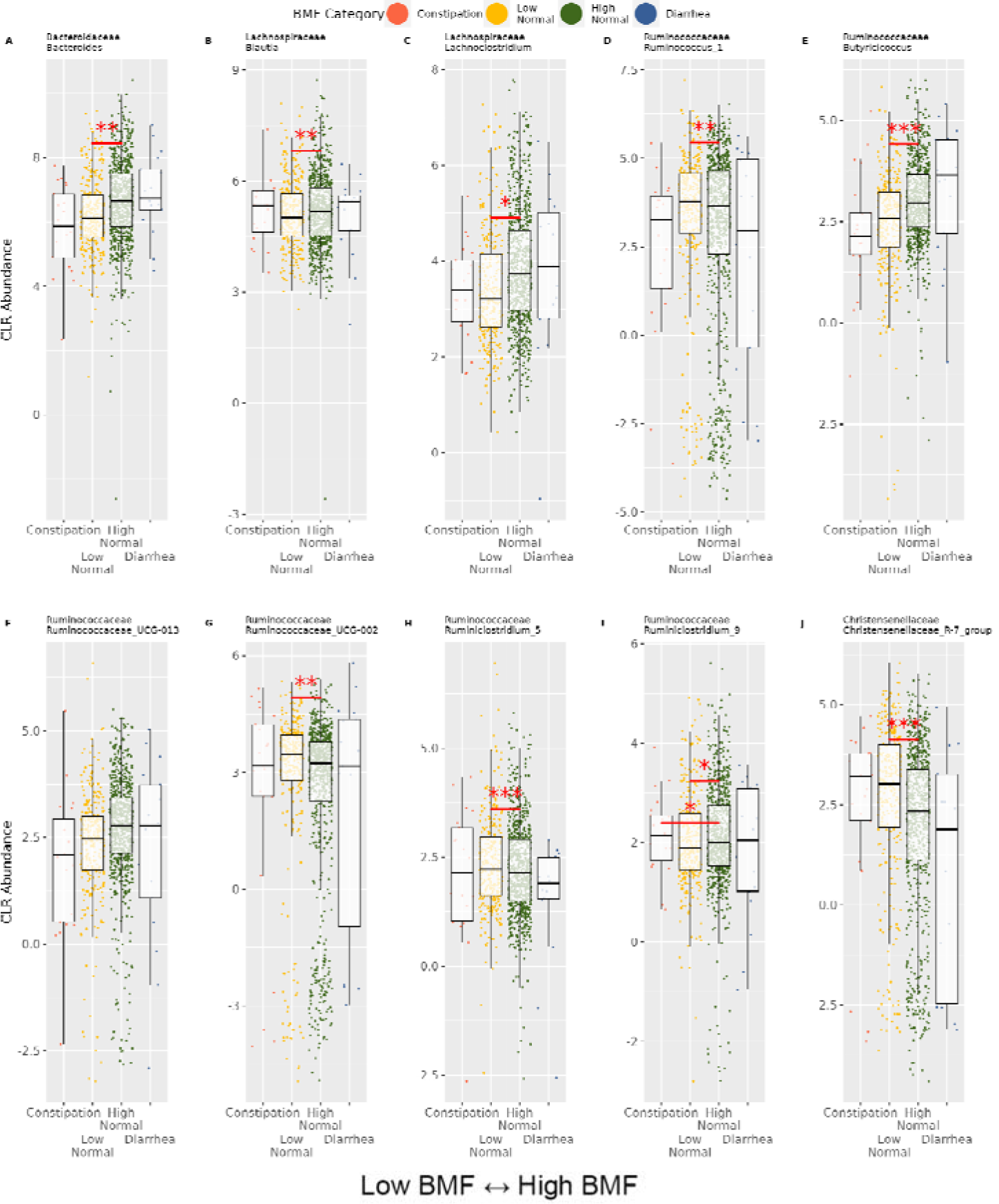
The top 10 most abundant genera significantly associated with BMF (A-J). Significant genera from the CORNCOB analysis in order of decreasing CLR-transformed abundance. The line in each plot denotes significant differences from the reference category (“High Normal” BMF), and asterisks denote FDR-corrected significance threshold. (***): p < 0.0001, (**): 0.0001 < p < 0.01, (*): 0.01 < p < 0.05. The horizontal axes are annotated as four BMF categories: “Constipation” (BMF = 1-2✕ per week), “Low Normal” (BMF = 3-6✕ per week), “High Normal” (BMF = 1-3✕ per day) which is the reference category in regression, and “Diarrhea” (BMF = 4✕ or more per day).

**Figure S3.**
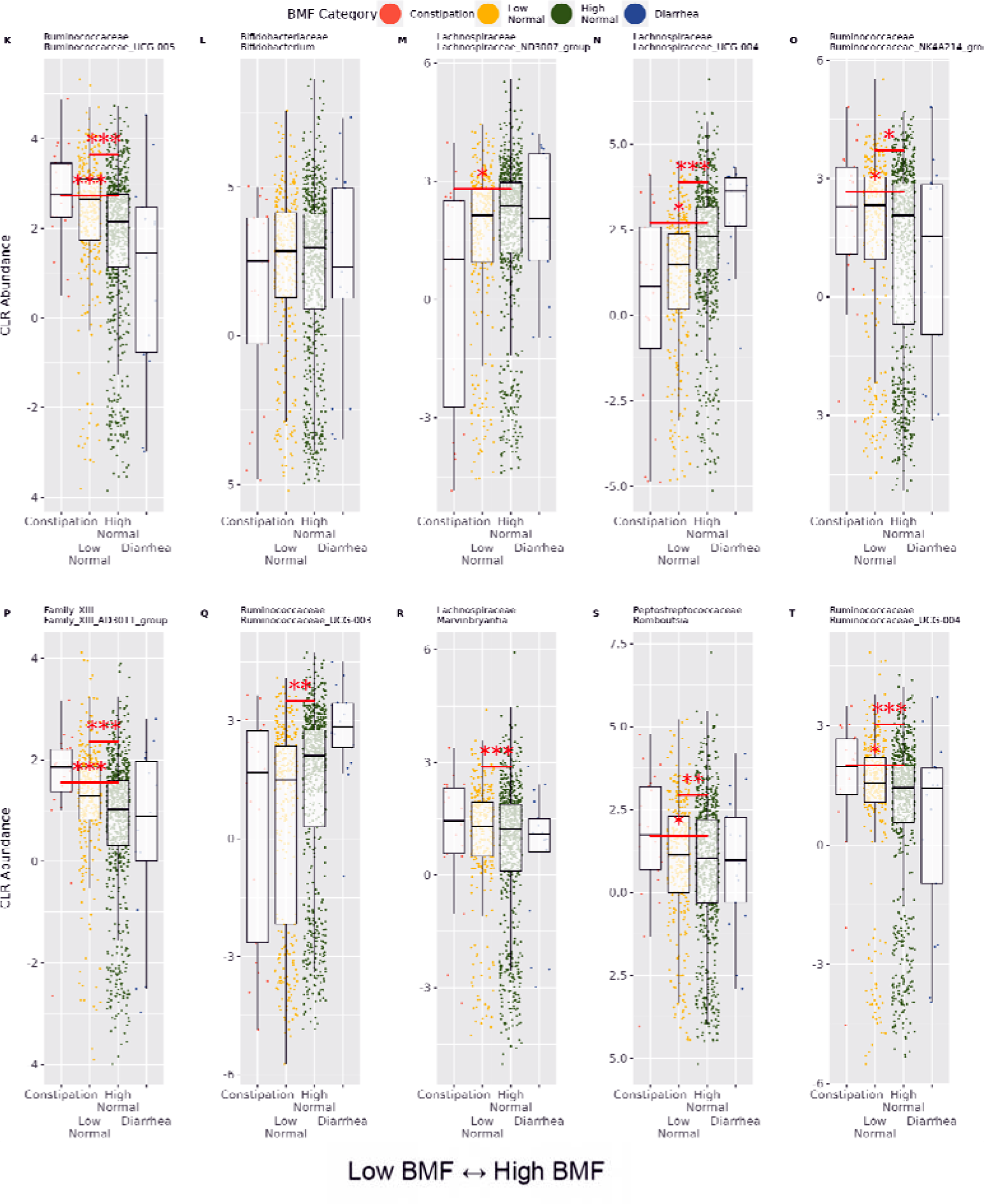
The top 11-20 most abundant genera associated with BMF (K-T). Significant genera from the CORNCOB analysis in order of decreasing CLR-transformed abundance. The line in each plot denotes significant differences from the reference category (“High Normal” BMF), and asterisks denote FDR-corrected significance threshold. (***): p < 0.0001, (**): 0.0001 < p < 0.01, (*): 0.01 < p < 0.05. The horizontal axes are annotated as four BMF categories: “Constipation” (BMF = 1-2✕ per week), “Low Normal” (BMF = 3-6✕ per week), “High Normal” (BMF = 1-3✕ per day) which is the reference category in regression, and “Diarrhea” (BMF = 4✕ or more per day). plasma metabolites from the LIMMA analysis. The horizontal axes are annotated as four BMF categories: “Constipation” (BMF = 1-2✕ per week), “Low Normal” (BMF = 3-6✕ per week), “High Normal” (BMF = 1-3✕ per day) which is the reference category in regression, and “Diarrhea” (BMF = 4✕ or more per day). Red significant comparison lines across each plot denote significant differences from the reference category (“High Normal” BMF), and asterisks denote FDR-corrected significance threshold. (***): p < 0.0001, (**): 0.0001 < p < 0.01, (*): 0.01 < p < 0.05.

**Figure S4.**
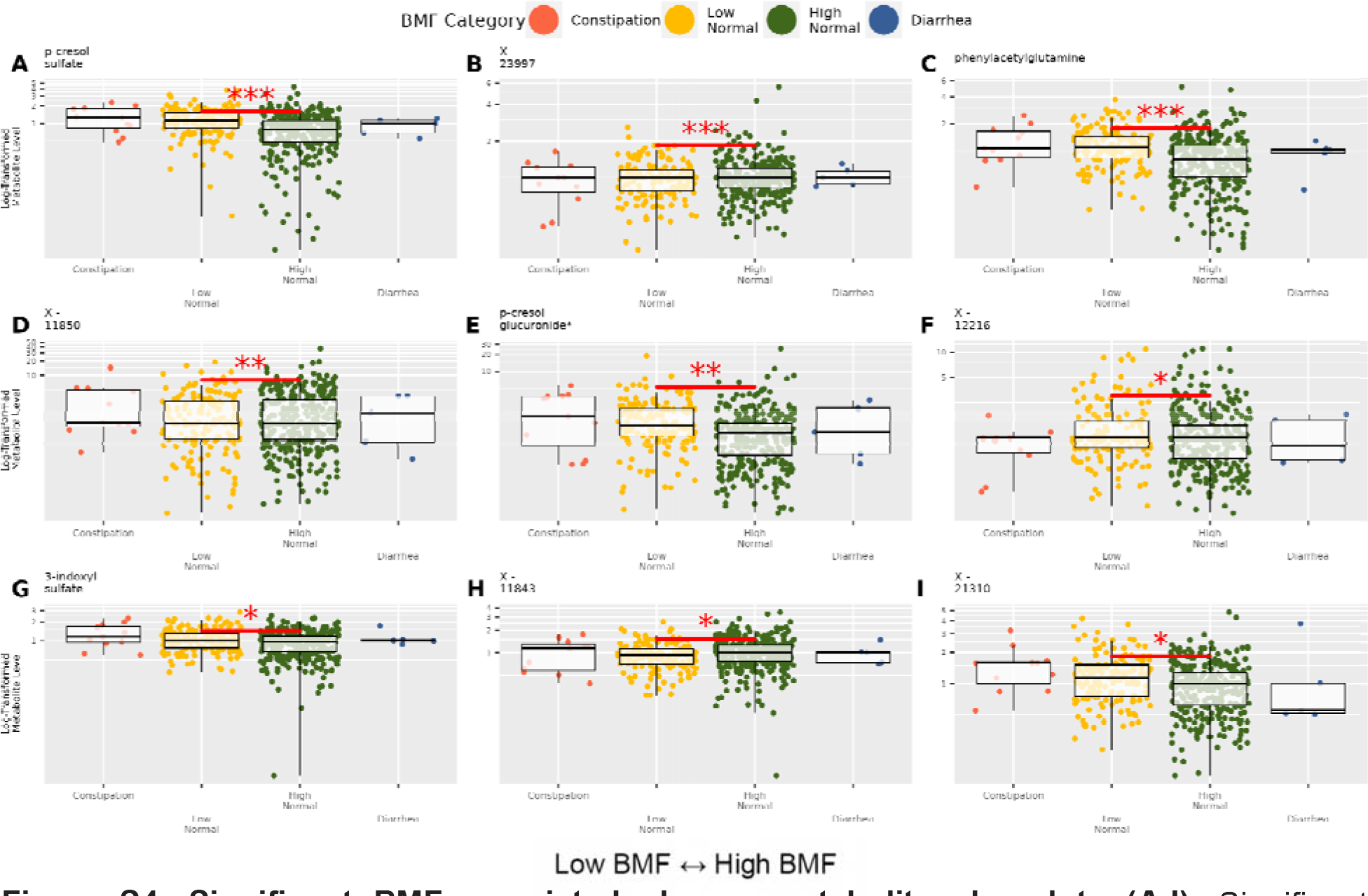
Significant BMF-associated pla sma metabolites boxplots (A-I). Significant plasma metabolites from the LIMMA analysis. The horizontal axes are annotated as four BMF categories: “Constipation” (BMF = 1-2✕ per week), “Low Normal” (BMF = 3-6✕ per week), “High Normal” (BMF = 1-3✕ per day) which is the reference category in regression, and “Diarrhea” (BMF = 4✕ or more per day). Red significant comparison lines across each plot denote significant differences from the reference category (“High Normal” BMF), and asterisks denote FDR-corrected significance threshold. (***): p < 0.0001, (**): 0.0001 < p < 0.01, (*): 0.01 < p < 0.05.

**Figure S5.**
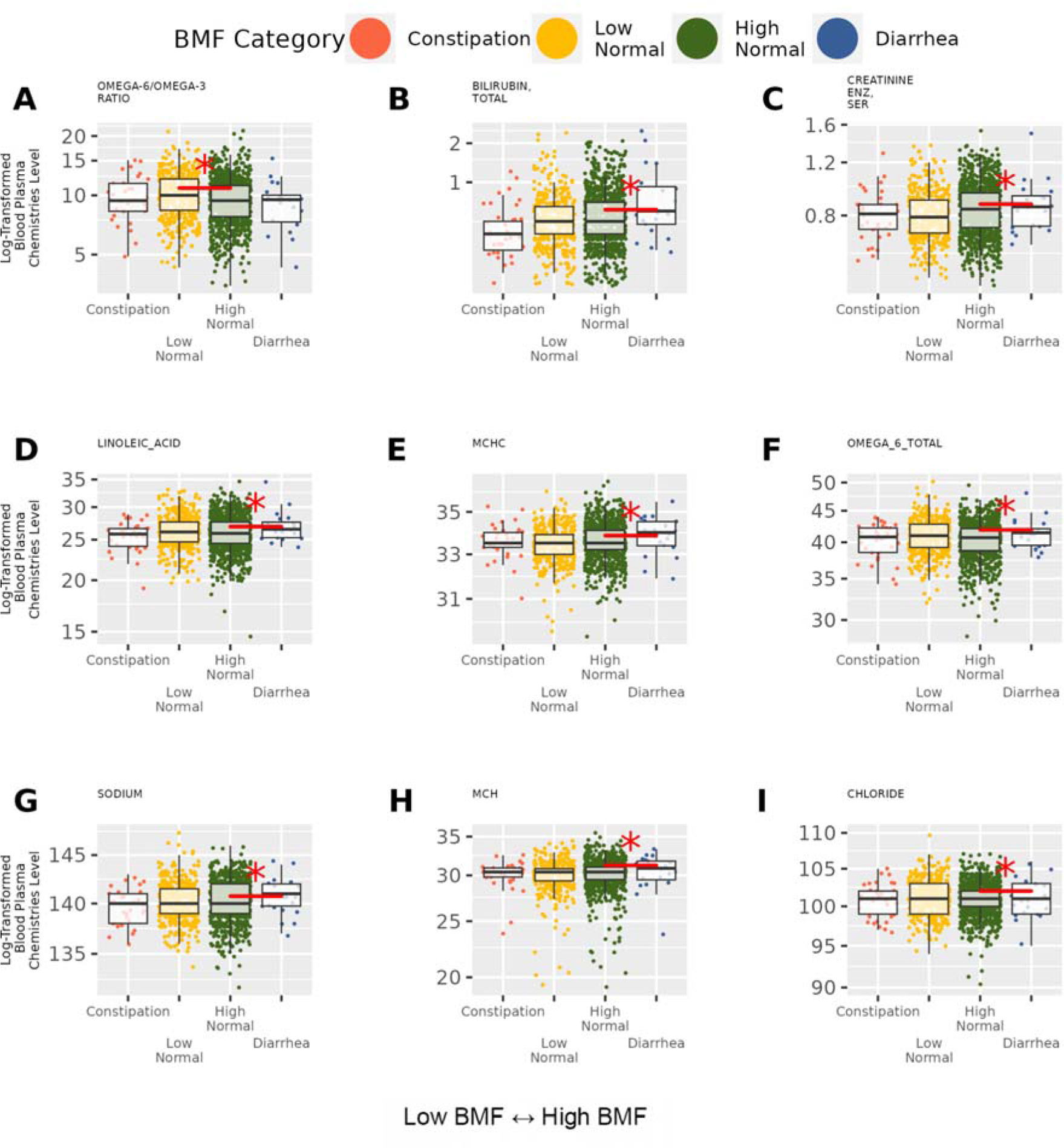
Significant BMF-associated clinical chemistries boxplots (A-I). Significant clinical chemistries from the LIMMA analysis. The horizontal axes are annotated as four BMF categories: “Constipation” (BMF = 1-2✕ per week), “Low Normal” (BMF = 3-6✕ per week), “High Normal” (BMF = 1-3✕ per day) which is the reference category in regression, and “Diarrhea” (BMF = 4✕ or more per day). Red significant comparison lines across each plot denote significant differences from the reference category (“High Normal” BMF), and asterisks denote FDR-corrected significance threshold. (***): p < 0.0001, (**): 0.0001 < p < 0.01, (*): 0.01 < p < 0.05.

**Figure S6.**
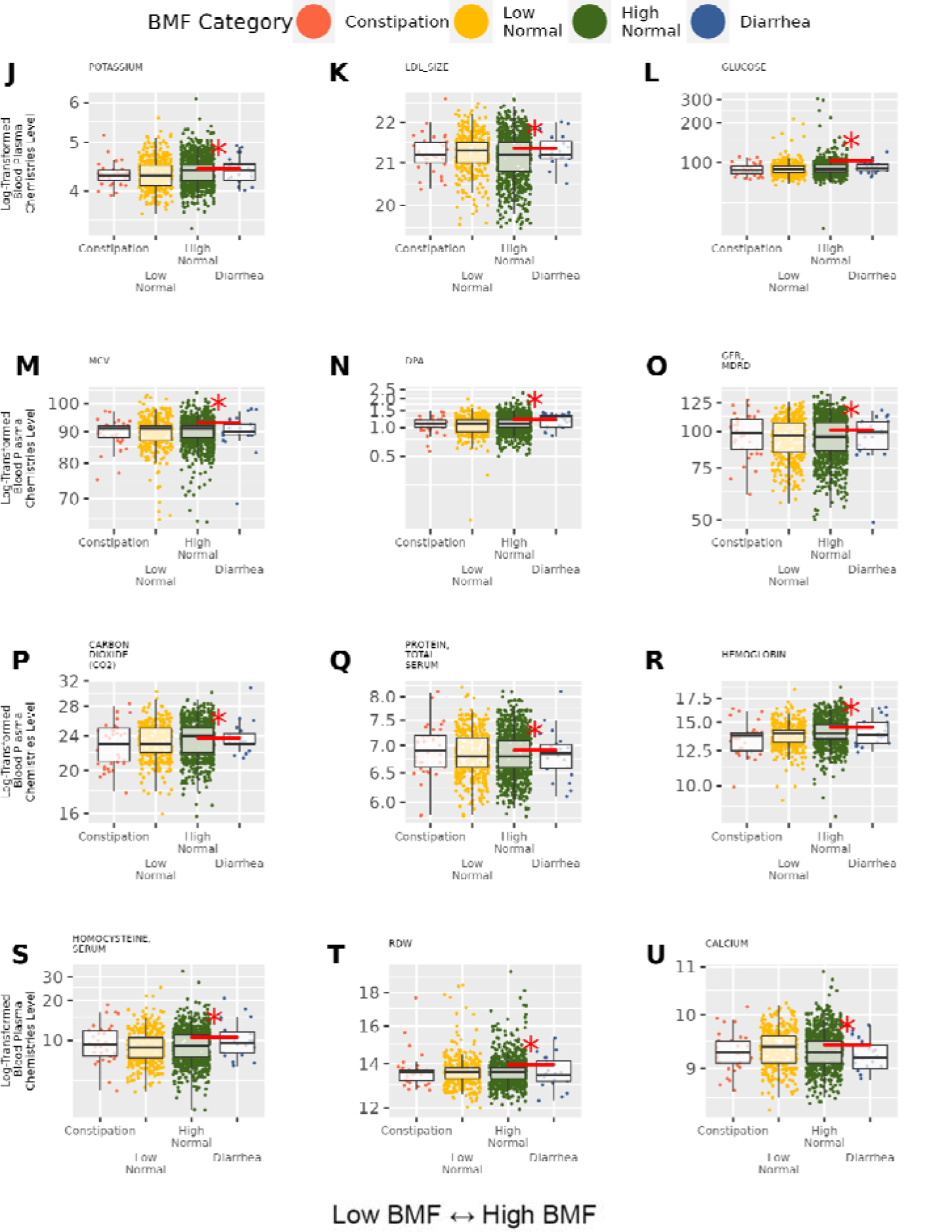
The remaining significant BMF-associated clinical chemistries boxplots (J-U). The remaining significant clinical chemistries from the LIMMA analysis. The horizontal axes are annotated as four BMF categories: “Constipation” (BMF = 1-2✕ per week), “Low Normal” (BMF = 3-6✕ per week), “High Normal” (BMF = 1-3✕ per day) which is the reference category in regression, and “Diarrhea” (BMF = 4✕ or more per day). Red significant comparison lines across each plot denote significant differences from the reference category (“High Normal” BMF), and asterisks denote FDR-corrected significance threshold. (***): p < 0.0001, (**): 0.0001 < p < 0.01, (*): 0.01 < p < 0.05.

**Figure S7.**
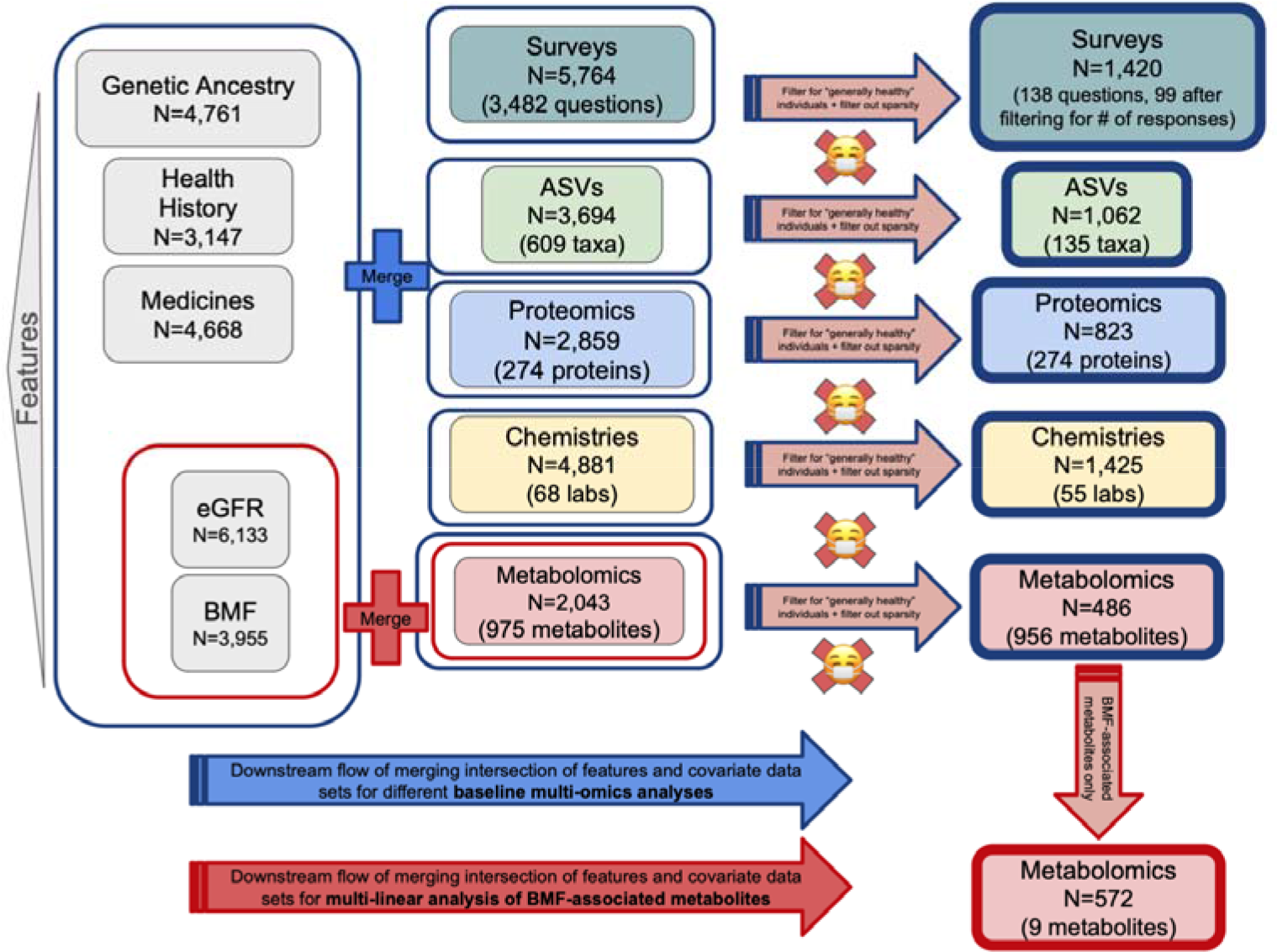
Flow Chart for Cohort Selection of Baseline Population. Individuals with the full complement of covariate data (sex, age, BMI, and CRP, LDL, A1C, and PCs1-3) were further filtered for having available baseline data for each of the following: surveys, microbiome profiles, proteomics, clinical chemistries (e.g. complete blood count, or CBC; and comprehensive metabolic panel, or CMP) and metabolomics. The “generally-healthy” exclusion criteria were then imposed (38.5% excluded; see Method Details), along with sparsity or non-missingness minimums for the features in the ‘omics data (≥ 30% prevalence for gut microbiome data, metabolomics and clinical chemistries; ≥ 50% prevalence for proteomics; and ≥ 90% prevalence and ≥ 10% affirmative for binary responses in the survey questions). These filters resulted in the final sub-cohort numbers shown on the right side of the figure in blue outlines. Additionally, the eGFR and BMF data frames were merged with the metabolomics data frame and filtered by the “generally-healthy” exclusionary criteria to achieve 572 participants with the data for the 9 BMF- associated metabolites eGFR regression and mediation analysis.

**Figure S8.**
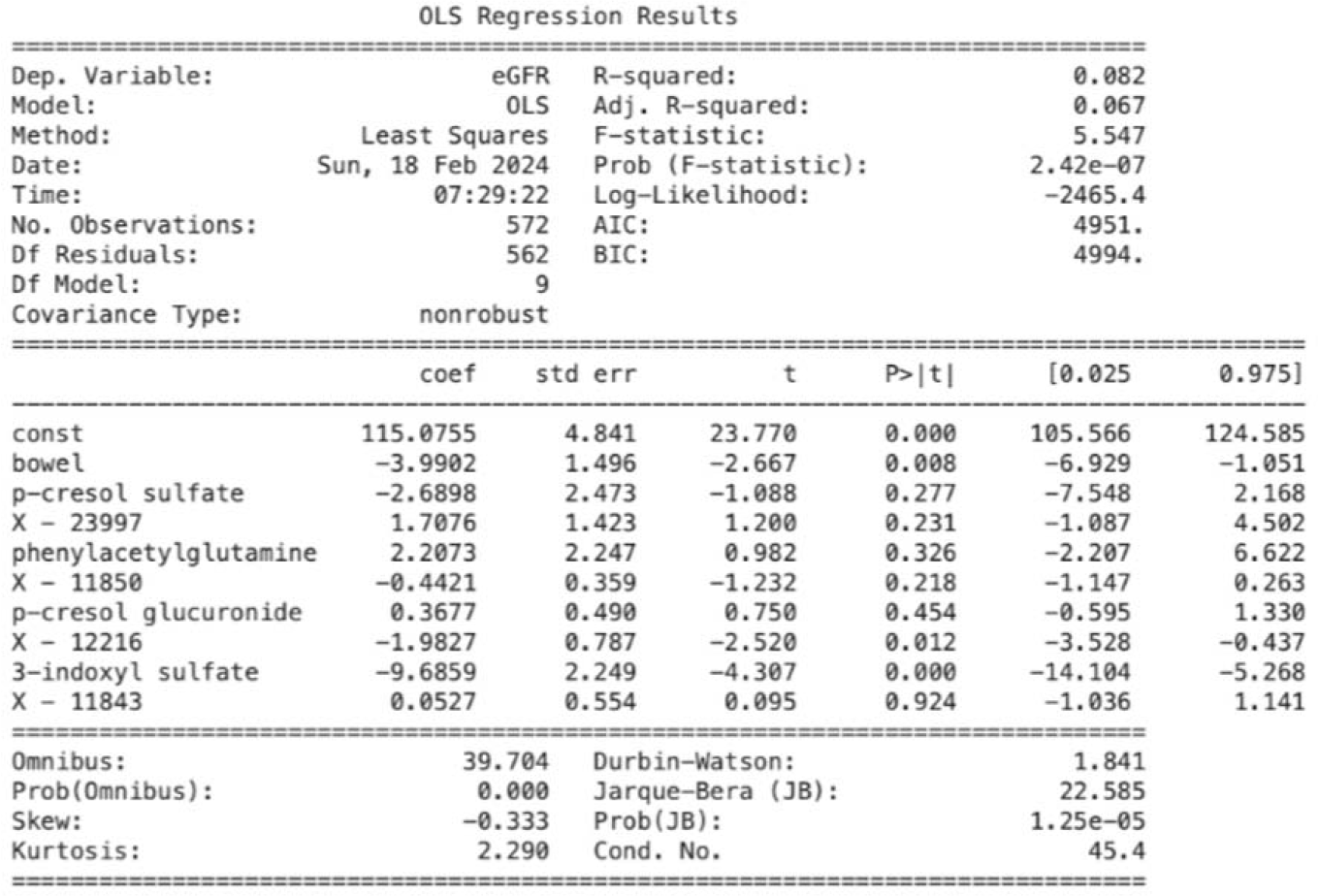
OLS regression resulting from eGFR ∼ BMF-associated metabolites + BMF. The p-value for the overall generalized-linear model (eGFR ∼ BMF-related metabolites) was significant (N = 572, p = 2.42E-7, R^2^ = 0.082) and so were the p-values of the individual β-coefficients for 3-IS (β_3-IS_ = -9.69, p = 1.96E-5), BMF (denoted “bowel”; β_BMF_ = -3.99, p = 7.88E- 3), and X - 12216 (β_X-12216_= -1.98, p = 1.20E-2).

## Notes

https://github.com/jajohnso29/Generally-Healthy-Cohort-BMF

### Summary of Updates

We stratified our initial population based on health status, and reran all of our analyses on a subset of generally-healthy individuals (i.e., no reported disease). We also include clinical chemistries and genetic ancestry as covariates in our regressions. Finally, we now find an association between a constipation-enriched metabolite in the blood and reduced kidney function (eGFR).

## REFERENCES

1. Magalhães-Guedes, K.T. (2022). Psychobiotic Therapy: Method to Reinforce the Immune System. Clin. Psychopharmacol. Neurosci. 20, 17–25. 10.9758/cpn.2022.20.1.17.

2. Kim, J., and Lee, H.K. (2021). Potential Role of the Gut Microbiome In Colorectal Cancer Progression. Front. Immunol. 12, 807648. 10.3389/fimmu.2021.807648.

3. Hughes, R.L. (2020). A Review of the Role of the Gut Microbiome in Personalized Sports Nutrition. Frontiers in Nutrition 6. 10.3389/fnut.2019.00191.

4. Ceballos, D., Hernández-Camba, A., and Ramos, L. (2021). Diet and microbiome in the beginning of the sequence of gut inflammation. World J Clin Cases 9, 11122–11147. 10.12998/wjcc.v9.i36.11122.

5. Cryan, J.F., O’Riordan, K.J., Cowan, C.S.M., Sandhu, K.V., Bastiaanssen, T.F.S., Boehme, M., Codagnone, M.G., Cussotto, S., Fulling, C., Golubeva, A.V., et al. (2019). The Microbiota-Gut-Brain Axis. Physiol. Rev. 99, 1877–2013. 10.1152/physrev.00018.2018.

6. Kim, D.S., Zhang, T., and Park, S. (2022). Protective effects of and water extract against memory deficits through the gut-microbiome-brain axis in an Alzheimer’s disease model. Pharm. Biol. 60, 212–224. 10.1080/13880209.2022.2025860.

7. 7. Asnicar, F., Leeming, E.R., Dimidi, E., Mazidi, M., Franks, P.W., Al Khatib, H., Valdes, A.M., Davies, R., Bakker, E., Francis, L., et al. (2021). Blue poo: impact of gut transit time on the gut microbiome using a novel marker. Gut 70, 1665–1674. 10.1136/gutjnl-2020-323877.

8. Müller, M., Canfora, E.E., and Blaak, E.E. (2018). Gastrointestinal Transit Time, Glucose Homeostasis and Metabolic Health: Modulation by Dietary Fibers. Nutrients 10. 10.3390/nu10030275.

9. Vanuytsel, T., Tack, J., and Farre, R. (2021). The Role of Intestinal Permeability in Gastrointestinal Disorders and Current Methods of Evaluation. Front Nutr 8, 717925. 10.3389/fnut.2021.717925.

10. Lewis, S.J., and Heaton, K.W. (1997). Stool form scale as a useful guide to intestinal transit time. Scand. J. Gastroenterol. 32, 920–924. 10.3109/00365529709011203.

11. Keendjele, T.P.T., Eelu, H.H., Nashihanga, T.E., Rennie, T.W., and Hunter, C.J. (2021). Corn? When did I eat corn? Gastrointestinal transit time in health science students. Adv. Physiol. Educ. 45, 103–108. 10.1152/advan.00192.2020.

12. Wilmanski, T., Rappaport, N., Earls, J.C., Magis, A.T., Manor, O., Lovejoy, J., Omenn, G.S., Hood, L., Gibbons, S.M., and Price, N.D. (2019). Blood metabolome predicts gut microbiome α-diversity in humans. Nature Biotechnology 37, 1217–1228. 10.1038/s41587-019-0233-9.

13. Saad, R.J., Rao, S.S.C., Koch, K.L., Kuo, B., Parkman, H.P., McCallum, R.W., Sitrin, M.D., Wilding, G.E., Semler, J.R., and Chey, W.D. (2010). Do stool form and frequency correlate with whole-gut and colonic transit? Results from a multicenter study in constipated individuals and healthy controls. Am. J. Gastroenterol. 105, 403–411. 10.1038/ajg.2009.612.

14. Roager, H.M., Hansen, L.B.S., Bahl, M.I., Frandsen, H.L., Carvalho, V., Gøbel, R.J., Dalgaard, M.D., Plichta, D.R., Sparholt, M.H., Vestergaard, H., et al. (2016). Colonic transit time is related to bacterial metabolism and mucosal turnover in the gut. Nature Microbiology 1, 1–9. 10.1038/nmicrobiol.2016.93.

15. Wiskur, B., and Meerveld, B.G.-V. (2010). The Aging Colon: The Role of Enteric Neurodegeneration in Constipation. Current Gastroenterology Reports 12, 507–512. 10.1007/s11894-010-0139-7.

16. Stirpe, P., Hoffman, M., Badiali, D., and Colosimo, C. (2016). Constipation: an emerging risk factor for Parkinson’s disease? European Journal of Neurology 23, 1606–1613. 10.1111/ene.13082.

17. Duvallet, C., Gibbons, S.M., Gurry, T., Irizarry, R.A., and Alm, E.J. (2017). Meta-analysis of gut microbiome studies identifies disease-specific and shared responses. Nat. Commun. 8, 1784. 10.1038/s41467-017-01973-8.

18. McDonald, D., Hyde, E., Debelius, J.W., Morton, J.T., Gonzalez, A., Ackermann, G., Aksenov, A.A., Behsaz, B., Brennan, C., Chen, Y., et al. (2018). American Gut: an Open Platform for Citizen Science Microbiome Research. mSystems 3. 10.1128/mSystems.00031-18.

19. Tomkovich, S., Taylor, A., King, J., Colovas, J., Bishop, L., McBride, K., Royzenblat, S., Lesniak, N.A., Bergin, I.L., and Schloss, P.D. (2021). An Osmotic Laxative Renders Mice Susceptible to Prolonged Clostridioides difficile Colonization and Hinders Clearance. mSphere 6. 10.1128/mSphere.00629-21.

20. Falony, G., Joossens, M., Vieira-Silva, S., Wang, J., Darzi, Y., Faust, K., Kurilshikov, A., Bonder, M.J., Valles-Colomer, M., Vandeputte, D., et al. (2016). Population-level analysis of gut microbiome variation. Science 352, 560–564. 10.1126/science.aad3503.

21. Vandeputte, D., Falony, G., Vieira-Silva, S., Tito, R.Y., Joossens, M., and Raes, J. (2016). Stool consistency is strongly associated with gut microbiota richness and composition, enterotypes and bacterial growth rates. Gut 65, 57–62. 10.1136/gutjnl-2015-309618.

22. Adams-Carr, K.L., Bestwick, J.P., Shribman, S., Lees, A., Schrag, A., and Noyce, A.J. (2016). Constipation preceding Parkinson’s disease: a systematic review and meta- analysis. J. Neurol. Neurosurg. Psychiatry 87, 710–716. 10.1136/jnnp-2015-311680.

23. Ramos, C.I., Nerbass, F.B., and Cuppari, L. (2022). Constipation in Chronic Kidney Disease: It Is Time to Bridge the Gap. Kidney and Dialysis 2, 221–233. 10.3390/kidneydial2020023.

24. Sumida, K., Yamagata, K., and Kovesdy, C.P. (2020). Constipation in CKD. Kidney Int Rep 5, 121–134. 10.1016/j.ekir.2019.11.002.

25. Ikee, R., Sasaki, N., Yasuda, T., and Fukazawa, S. (2020). Chronic Kidney Disease, Gut Dysbiosis, and Constipation: A Burdensome Triplet. Microorganisms 8, 1862. 10.3390/microorganisms8121862.

26. Gr, B.A.P.J. (2013). American Gastroenterological Association technical review on constipation. Gastroenterology 144, 218–238. 10.1053/j.gastro.2012.10.028. .

27. Sumida, K., Molnar, M.Z., Potukuchi, P.K., Thomas, F., Lu, J.L., Matsushita, K., Yamagata, K., Kalantar-Zadeh, K., and Kovesdy, C.P. (2017). Constipation and Incident CKD. Journal of the American Society of Nephrology 28, 1248–1258. 10.1681/asn.2016060656.

28. 28. Brocca, A., Virzì, G.M., de Cal, M., Cantaluppi, V., and Ronco, C. (2013). Cytotoxic effects of p-cresol in renal epithelial tubular cells. Blood Purif. 36, 219–225. 10.1159/000356370.

29. 29. Poesen, R., Claes, K., Evenepoel, P., de Loor, H., Augustijns, P., Kuypers, D., and Meijers, B. (2016). Microbiota-Derived Phenylacetylglutamine Associates with Overall Mortality and Cardiovascular Disease in Patients with CKD. J. Am. Soc. Nephrol. 27, 3479–3487. 10.1681/ASN.2015121302.

30. Barrios, C., Beaumont, M., Pallister, T., Villar, J., Goodrich, J.K., Clark, A., Pascual, J., Ley, R.E., Spector, T.D., Bell, J.T., et al. (2015). Gut-Microbiota-Metabolite Axis in Early Renal Function Decline. PLoS One 10, e0134311. 10.1371/journal.pone.0134311.

31. Ikee, R., Yano, K., and Tsuru, T. (2019). Constipation in chronic kidney disease: it is time to reconsider. Renal Replacement Therapy 5. 10.1186/s41100-019-0246-3.

32. Martin, B.D., Witten, D., and Willis, A.D. (2020). MODELING MICROBIAL ABUNDANCES AND DYSBIOSIS WITH BETA-BINOMIAL REGRESSION. Ann. Appl. Stat. 14, 94–115. 10.1214/19-aoas1283.

33. polr function - RDocumentation https://www.rdocumentation.org/packages/MASS/versions/7.3-58.1/topics/polr.

34. [No title] https://cran.r-project.org/web/packages/mediation/vignettes/mediation.pdf.

35. Wan, M., King, L., Baugh, N., Arslan, Z., Snauwaert, E., Paglialonga, F., and Shroff, R. (2023). Gutted: constipation in children with chronic kidney disease and on dialysis. Pediatr. Nephrol. 10.1007/s00467-022-05849-y.

36. Kim, J.E., Park, J.J., Lee, M.R., Choi, J.Y., Song, B.R., Park, J.W., Kang, M.J., Son, H.J., Hong, J.T., and Hwang, D.Y. (2019). Constipation in Tg2576 mice model for Alzheimer’s disease associated with dysregulation of mechanism involving the mAChR signaling pathway and ER stress response. PLoS One 14, e0215205. 10.1371/journal.pone.0215205.

37. Baldini, F., Hertel, J., Sandt, E., Thinnes, C.C., Neuberger-Castillo, L., Pavelka, L., Betsou, F., Krüger, R., Thiele, I., and NCER-PD Consortium (2020). Parkinson’s disease- associated alterations of the gut microbiome predict disease-relevant changes in metabolic functions. BMC Biol. 18, 62. 10.1186/s12915-020-00775-7.

38. Neugarten, J., and Golestaneh, L. (2013). Gender and the Prevalence and Progression of Renal Disease. Advances in Chronic Kidney Disease 20, 390–395. 10.1053/j.ackd.2013.05.004.

39. Werth, B.L., and Christopher, S.-A. (2021). Potential risk factors for constipation in the community. World J. Gastroenterol. 27, 2795–2817. 10.3748/wjg.v27.i21.2795.

40. Chen, H.B., Huang, Y., Song, H.W., Li, X.L., He, S., Xie, J.T., Huang, C., Zhang, S.J., Liu, J., and Zou, Y. (2010). Clinical Research on the Relation Between Body Mass Index, Motilin and Slow Transit Constipation. Gastroenterol. Res. Pract. 3, 19–24. 10.4021/gr2010.02.168w.

41. Vermorken, A.J.M., Andrès, E., and Cui, Y. (2016). Bowel movement frequency, oxidative stress and disease prevention. Mol Clin Oncol 5, 339–342. 10.3892/mco.2016.987.

42. Vermorken, A.J.M., Cui, Y., Kleerebezem, R., and Andrès, E. (2016). Bowel movement frequency and cardiovascular mortality, a matter of fibers and oxidative stress? Atherosclerosis 253, 278–280. 10.1016/j.atherosclerosis.2016.08.012.

43. Jankipersadsing, S.A., Hadizadeh, F., Bonder, M.J., Tigchelaar, E.F., Deelen, P., Fu, J., Andreasson, A., Agreus, L., Walter, S., Wijmenga, C., et al. (2017). A GWAS meta-analysis suggests roles for xenobiotic metabolism and ion channel activity in the biology of stool frequency. Gut 66, 756–758. 10.1136/gutjnl-2016-312398.

44. Romano, S., Savva, G.M., Bedarf, J.R., Charles, I.G., Hildebrand, F., and Narbad, A. (2021). Meta-analysis of the Parkinson’s disease gut microbiome suggests alterations linked to intestinal inflammation. NPJ Parkinson’s Disease 7. 10.1038/s41531-021-00156-z.

45. Singh, S.B., Carroll-Portillo, A., and Lin, H.C. (2023). Desulfovibrio in the Gut: The Enemy within? Microorganisms 11. 10.3390/microorganisms11071772.

46. Boutard, M., Cerisy, T., Nogue, P.-Y., Alberti, A., Weissenbach, J., Salanoubat, M., and Tolonen, A.C. (2014). Functional Diversity of Carbohydrate-Active Enzymes Enabling a Bacterium to Ferment Plant Biomass. PLoS Genet. 10. 10.1371/journal.pgen.1004773.

47. Song, Y., Chen, K., Lv, L., Xiang, Y., Du, X., Zhang, X., Zhao, G., and Xiao, Y. (2022). Uncovering the biogeography of the microbial commmunity and its association with nutrient metabolism in the intestinal tract using a pig model. Front. Nutr. 9, 1003763. 10.3389/fnut.2022.1003763.

48. 48. van den Bogert, B., Meijerink, M., Zoetendal, E.G., Wells, J.M., and Kleerebezem, M. (2014). Immunomodulatory Properties of Streptococcus and Veillonella Isolates from the Human Small Intestine Microbiota. PLoS One 9, e114277. 10.1371/journal.pone.0114277.

49. Silva, Y.P., Bernardi, A., and Frozza, R.L. (2020). The Role of Short-Chain Fatty Acids From Gut Microbiota in Gut-Brain Communication. Front. Endocrinol. 11, 508738. 10.3389/fendo.2020.00025.

50. Martin-Gallausiaux, C., Marinelli, L., Blottière, H.M., Larraufie, P., and Lapaque, N. (2021). SCFA: mechanisms and functional importance in the gut. Proc. Nutr. Soc. 80, 37–49. 10.1017/S0029665120006916.

51. Zhao, Y., Liu, Q., Hou, Y., and Zhao, Y. (2022). Alleviating effects of gut micro-ecologically regulatory treatments on mice with constipation. Front. Microbiol. 13, 956438. 10.3389/fmicb.2022.956438.

52. Vicentini, F.A., Keenan, C.M., Wallace, L.E., Woods, C., Cavin, J.-B., Flockton, A.R., Macklin, W.B., Belkind-Gerson, J., Hirota, S.A., and Sharkey, K.A. (2021). Intestinal microbiota shapes gut physiology and regulates enteric neurons and glia. Microbiome 9, 210. 10.1186/s40168-021-01165-z.

53. Bodnar, H.B., and Bodnar, B.M. (2014). [Peculiarities of a large bowel microstructure in children, suffering chronic constipation, caused by its inborn anomaly]. Klin. Khir., 30–33.

54. Xiao, Y., and Kashyap, P.C. (2022). Microbially derived polyunsaturated fatty acid as a modulator of gastrointestinal motility. J. Clin. Invest. 132. 10.1172/JCI161572.

55. Negussie, A.B., Dell, A.C., Davis, B.A., and Geibel, J.P. (2022). Colonic Fluid and Electrolyte Transport 2022: An Update. Cells 11. 10.3390/cells11101712.

56. Clayburgh, D.R., Shen, L., and Turner, J.R. (2004). A porous defense: the leaky epithelial barrier in intestinal disease. Lab. Invest. 84, 282–291. 10.1038/labinvest.3700050.

57. Mitsuoka, T. (1996). Intestinal flora and human health. Asia Pac. J. Clin. Nutr. 5, 2–9.

58. Usuda, H., Okamoto, T., and Wada, K. (2021). Leaky Gut: Effect of Dietary Fiber and Fats on Microbiome and Intestinal Barrier. Int. J. Mol. Sci. 22. 10.3390/ijms22147613.

59. Zhang, L., Xie, F., Tang, H., Zhang, X., Hu, J., Zhong, X., Gong, N., Lai, Y., Zhou, M., Tian, J., et al. (2022). Gut microbial metabolite TMAO increases peritoneal inflammation and peritonitis risk in peritoneal dialysis patients. Transl. Res. 240, 50–63. 10.1016/j.trsl.2021.10.001.

60. Yu, W., Venkatraman, A., Menden, H.L., Martinez, M., Umar, S., and Sampath, V. (2023). Short-chain fatty acids ameliorate necrotizing enterocolitis-like intestinal injury through enhancing Notch1-mediated single immunoglobulin interleukin-1-related receptor, toll- interacting protein, and A20 induction. Am. J. Physiol. Gastrointest. Liver Physiol. 324, G24–G37. 10.1152/ajpgi.00057.2022.

61. Chen, Y.-Y., Chen, D.-Q., Chen, L., Liu, J.-R., Vaziri, N.D., Guo, Y., and Zhao, Y.-Y. (2019). Microbiome–metabolome reveals the contribution of gut–kidney axis on kidney disease. J. Transl. Med. 17, 1–11. 10.1186/s12967-018-1756-4.

62. 62. Hobson, S., Qureshi, A.R., Ripswedan, J., Wennberg, L., de Loor, H., Ebert, T., Söderberg, M., Evenepoel, P., Stenvinkel, P., and Kublickiene, K. (2023). Phenylacetylglutamine and trimethylamine N-oxide: Two uremic players, different actions. Eur. J. Clin. Invest. 53, e14074. 10.1111/eci.14074.

63. Sun, C.-Y., Li, J.-R., Wang, Y.-Y., Lin, S.-Y., Ou, Y.-C., Lin, C.-J., Wang, J.-D., Liao, S.-L., and Chen, C.-J. (2020). p-Cresol Sulfate Caused Behavior Disorders and Neurodegeneration in Mice with Unilateral Nephrectomy Involving Oxidative Stress and Neuroinflammation. Int. J. Mol. Sci. 21. 10.3390/ijms21186687.

64. Levi, I., Gurevich, M., Perlman, G., Magalashvili, D., Menascu, S., Bar, N., Godneva, A., Zahavi, L., Chermon, D., Kosower, N., et al. (2021). Potential role of indolelactate and butyrate in multiple sclerosis revealed by integrated microbiome-metabolome analysis. Cell Rep Med 2, 100246. 10.1016/j.xcrm.2021.100246.

65. Sun, C.-Y., Hsu, H.-H., and Wu, M.-S. (2012). p-Cresol sulfate and indoxyl sulfate induce similar cellular inflammatory gene expressions in cultured proximal renal tubular cells. Nephrol. Dial. Transplant 28, 70–78. 10.1093/ndt/gfs133.

66. Barreto, F.C., Barreto, D.V., Liabeuf, S., Meert, N., Glorieux, G., Temmar, M., Choukroun, G., Vanholder, R., Massy, Z.A., and European Uremic Toxin Work Group (EUTox) (2009). Serum indoxyl sulfate is associated with vascular disease and mortality in chronic kidney disease patients. Clin. J. Am. Soc. Nephrol. 4, 1551–1558. 10.2215/CJN.03980609.

67. Tsai, M.-T., and Tarng, D.-C. (2018). Beyond a Measure of Liver Function—Bilirubin Acts as a Potential Cardiovascular Protector in Chronic Kidney Disease Patients. Int. J. Mol. Sci. 20, 117. 10.3390/ijms20010117.

68. 68. Creatinine test (2023). https://www.mayoclinic.org/tests-procedures/creatinine-test/about/pac-20384646.

69. 69. Creatinine (2023). National Kidney Foundation. https://www.kidney.org/atoz/content/serum-blood-creatinine.

70. 70. Llc, H. io Linoleic Acid. https://healthmatters.io/understand-blood-test-results/linoleic-acid.

71. 71. Sherrell, Z., and Labedzki, M. (2022). What are normal MCH levels? The Checkup. https://www.singlecare.com/blog/mch-blood-test/.

72. Hosseinzadeh, S.T., Poorsaadati, S., Radkani, B., and Forootan, M. (2011). Psychological disorders in patients with chronic constipation. Gastroenterol Hepatol Bed Bench 4, 159– 163.

73. 73. Blanc, F., Bouteloup, V., Paquet, C., Chupin, M., Pasquier, F., Gabelle, A., Ceccaldi, M., de Sousa, P.L., Krolak-Salmon, P., David, R., et al. (2022). Prodromal characteristics of dementia with Lewy bodies: baseline results of the MEMENTO memory clinics nationwide cohort. Alzheimers. Res. Ther. 14, 96. 10.1186/s13195-022-01037-0.

74. Nedelec, T., Couvy-Duchesne, B., Monnet, F., Daly, T., Ansart, M., Gantzer, L., Lekens, B., Epelbaum, S., Dufouil, C., and Durrleman, S. (2022). Identifying health conditions associated with Alzheimer’s disease up to 15 years before diagnosis: an agnostic study of French and British health records. Lancet Digit Health 4, e169–e178. 10.1016/S2589-7500(21)00275-2.

75. Kuang, R., and Binion, D.G. (2022). Should high-fiber diets be recommended for patients with inflammatory bowel disease? Curr. Opin. Gastroenterol. 38, 168–172. 10.1097/MOG.0000000000000810.

76. 76. Eating right for chronic kidney disease (2022). National Institute of Diabetes and Digestive and Kidney Diseases. https://www.niddk.nih.gov/health-information/kidney-disease/chronic-kidney-disease-ckd/eating-nutrition.

77. O’Rourke, H.P., and MacKinnon, D.P. (2018). Reasons for Testing Mediation in the Absence of an Intervention Effect: A Research Imperative in Prevention and Intervention Research. J. Stud. Alcohol Drugs 79, 171. 10.15288/jsad.2018.79.171.

78. Frequent Bowel Movements Cleveland Clinic. https://my.clevelandclinic.org/health/diseases/17791-frequent-bowel-movements.

79. Callahan, B.J., McMurdie, P.J., Rosen, M.J., Han, A.W., Johnson, A.J.A., and Holmes, S.P. (2016). DADA2: High-resolution sample inference from Illumina amplicon data. Nat. Methods 13, 581–583. 10.1038/nmeth.3869.

80. Quast, C., Pruesse, E., Yilmaz, P., Gerken, J., Schweer, T., Yarza, P., Peplies, J., and Glöckner, F.O. (2013). The SILVA ribosomal RNA gene database project: improved data processing and web-based tools. Nucleic Acids Res. 41, D590–D596. 10.1093/nar/gks1219.

81. McMurdie, P.J., and Holmes, S. (2013). phyloseq: an R package for reproducible interactive analysis and graphics of microbiome census data. PLoS One 8, e61217. 10.1371/journal.pone.0061217.

82. [No title] https://www.illumina.com/content/dam/illumina/gcs/assembled-assets/marketing-literature/olink-proteomics-tech-note-m-us-00196/olink-proteomics-tech-note-m-us-00196.pdf.

83. 83. Assay validation (2021). Olink. https://olink.com/our-platform/assay-validation/.

84. Wik, L., Nordberg, N., Broberg, J., Björkesten, J., Assarsson, E., Henriksson, S., Grundberg, I., Pettersson, E., Westerberg, C., Liljeroth, E., et al. (2021). Proximity Extension Assay in Combination with Next-Generation Sequencing for High-throughput Proteome-wide Analysis. Mol. Cell. Proteomics 20, 100168. 10.1016/j.mcpro.2021.100168.

85. Zubair, N., Conomos, M.P., Hood, L., Omenn, G.S., Price, N.D., Spring, B.J., Magis, A.T., and Lovejoy, J.C. (2019). Genetic Predisposition Impacts Clinical Changes in a Lifestyle Coaching Program. Sci. Rep. 9. 10.1038/s41598-019-43058-0.

86. Manor, O., Zubair, N., Conomos, M.P., Xu, X., Rohwer, J.E., Krafft, C.E., Lovejoy, J.C., and Magis, A.T. (2018). A Multi-omic Association Study of Trimethylamine N-Oxide. Cell Rep. 24, 935–946. 10.1016/j.celrep.2018.06.096.

87. Diener, C., Dai, C.L., Wilmanski, T., Baloni, P., Smith, B., Rappaport, N., Hood, L., Magis, A.T., and Gibbons, S.M. (2022). Genome-microbiome interplay provides insight into the determinants of the human blood metabolome. Nat Metab 4, 1560–1572. 10.1038/s42255-022-00670-1.

88. [No title] https://cran.r-project.org/web/packages/mediation/vignettes/mediation.pdf.

89. Delgado, C., Baweja, M., Crews, D.C., Eneanya, N.D., Gadegbeku, C.A., Inker, L.A., Mendu, M.L., Miller, W.G., Moxey-Mims, M.M., Roberts, G.V., et al. (2022). A Unifying Approach for GFR Estimation: Recommendations of the NKF-ASN Task Force on Reassessing the Inclusion of Race in Diagnosing Kidney Disease. Am. J. Kidney Dis. 79, 268–288.e1. 10.1053/j.ajkd.2021.08.003.

90. Ritchie, M.E., Phipson, B., Wu, D., Hu, Y., Law, C.W., Shi, W., and Smyth, G.K. (2015). limma powers differential expression analyses for RNA-sequencing and microarray studies. Nucleic Acids Res. 43, e47–e47. 10.1093/nar/gkv007.

